# Efficiently summarizing relationships in large samples: a general duality between statistics of genealogies and genomes

**DOI:** 10.1101/779132

**Authors:** Peter Ralph, Kevin Thornton, Jerome Kelleher

## Abstract

As a genetic mutation is passed down across generations, it distinguishes those genomes that have inherited it from those that have not, providing a glimpse of the genealogical tree relating the genomes to each other at that site. Statistical summaries of genetic variation therefore also describe the underlying genealogies. We use this correspondence to define a general framework that efficiently computes single-site population genetic statistics using the succinct tree sequence encoding of genealogies and genome sequence. The general approach accumulates “sample weights” within the genealogical tree at each position on the genome, which are then combined using a “summary function”; different statistics result from different choices of weight and function. Results can be reported in three ways: by *site,* which corresponds to statistics calculated as usual from genome sequence; by *branch,* which gives the expected value of the dual site statistic under the infinite-sites model of mutation, and by *node,* which summarizes the contribution of each ancestor to these statistics. We use the framework to implement many currently-defined statistics of genome sequence (making the statistics’ relationship to the underlying genealogical trees concrete and explicit), as well as the corresponding “branch” statistics of tree shape. We evaluate computational performance using simulated data, and show that calculating statistics from tree sequences using this general framework is several orders of magnitude more efficient than optimized matrix-based methods in terms of both run time and memory requirements. We also explore how well the duality between site and branch statistics holds in practice on trees inferred from the 1000 Genomes Project dataset, and discuss ways in which deviations may encode interesting biological signals.

## Introduction

It was once a major undertaking to collect data sufficient to estimate a single summary of the genetic relationships between the individuals of a sample [e.g., Kreitman, 1983]. Today’s vast quantity of wholegenome sequence makes it possible to confidently estimate many more properties of genealogical relationships in local regions of genomes, both between individuals [e.g., Browning and Browning, 2010, Aguillon et al., 2017] and within populations [e.g., Booker and Keightley, 2018, Haenel et al., 2018, Stankowski et al., 2019]. Computation is beginning to be a major problem: projects such as UK Biobank [Bycroft et al., 2018] and gnomAD [Karczewski et al., 2019] hold genetic data for hundreds of thousands of samples at tens to hundreds of millions of variant sites. Such large genotype matrices are extremely unwieldy and difficult to process, and computational complexity is usually linear in the size of the matrix. Tools such as `Hail` (https://hail.is) allow computations to be sharded across many machines in parallel, thereby calculating statistics far more quickly than is possible using single-computer methods such as `plink` [Purcell et al., 2007]. Computer time must still be paid for, however, and running calculations concurrently on thousands of cores quickly becomes expensive. Larger datasets still, consisting of millions of whole genomes, are currently being collected and it is clear that processing and storing these genotype matrices will be very costly.

Genotype matrices are massively redundant, however, and several specialized compression methods have been proposed [Christley et al., 2008, Qiao et al., 2012, Durbin, 2014, Sambo et al., 2014, Layer et al., 2016, Danek and Deorowicz, 2018, Lin et al., 2019]. The fundamental reason for this redundancy is that samples share ancestry: related individuals will tend to share the same state at a particular variant site. Trees describe these ancestral relationships [Felsenstein, 2004, Semple and Steel, 2003], and provide a natural and elegant way of encoding genetic data, not only recording the history of a sample but also massively compressing the genotype matrix [Ané and Sanderson, 2005]. Until recently, however, tree based approaches could not be used to encode data from sexually reproducing organisms, because recombination results in many trees along the genome [Hudson, 1983, Griffiths, 1991, Griffiths and Marjoram, 1996, Rasmussen et al., 2014], and there was no efficient way of representing this information. The *succinct tree sequence* (or *tree sequence,* for brevity) is a recently introduced data structure that encodes these genealogical trees along a genome concisely. It was introduced in the context of coalescent simulation [Kelleher et al., 2016], leading to scalability increases of several orders of magnitude over existing methods. The methods were subsequently extended and refined for forwards-time simulations [Kelleher et al., 2018, Haller et al., 2018], with similarly large efficiency gains. Recent work has shown that tree sequence algorithms can also be used to massively increase the scalability of methods for inferring genome-wide genealogies, and making it possible to infer trees for millions of samples [Kelleher et al., 2019]. The key to the remarkable efficiency of tree sequence algorithms is the way that shared structure in adjacent trees along the genome is encoded. As one looks across the genome, genealogies change at recombination breakpoints in ancestors. However, nearby trees tend to share much of their structure, and single genealogical relationships (e.g., individual *x* inherits from individual *y*) are often shared across relatively long distances, which manifest as shared *edges* across many adjacent trees in the tree sequence. This redundancy can be exploited to store genome sequence and the associated genealogical relationships very compactly, and formed the basis for efficient algorithms to calculate several population genetics statistics [Kelleher et al., 2016]. This paper generalizes the approach of Kelleher et al. [2016] to a much broader class of statistics, supporting arbitrary tree topologies and patterns of allelic variation, including polyallelic sites and back mutations.

In this paper, we study “single-site” genetic statistics, i.e., statistics of aligned genome sequence that can be expressed as averages over values computed separately for each site. We develop a theoretical and computational framework that encompasses a large class of population genetic statistics, generalizing many classical summaries of genetic variation. We define and explore a duality between “Site” and “Branch” statistics, and provide a detailed description (and analysis) of an efficient algorithm to compute statistics in this general setting. The methods we describe are implemented, tested, and validated in the `tskit` Python and C libraries, freely available at https://github.com/tskit-dev/tskit under the terms of the MIT license.

The basic relationship between tree shape and summaries of genetic variation that we will explain and utilize below is this: if mutations are neutral, then the expected number that occur on a given branch of a tree is proportional to the length of that branch. This “duality” (we will use the term more precisely later) has been used extensively in deriving properties of statistics in models of random mating [e.g., Tajima, 1983, Tavarée, 1984, Fu, 1995], as has the fact that it applies regardless of the underlying demographic model [e.g., Gillespie and Langley, 1979, Hudson, 1983, Slatkin, 1991, McVean, 2002, Lohse et al., 2016, Ralph, 2019]. We build on this central insight to define a flexible and computationally efficient framework for population genetic statistics on tree sequences. Roughly speaking, to compute a statistic we will propagate “weights” additively up each marginal tree, summarize these weights on each branch with a “summary function”, and aggregate these summaries to produce statistics. Different choices of weights and summary functions will then produce both familiar and novel statistics of both genotypes and genealogies.

## Framework and statistics

Population genetic data is most directly represented as a matrix of genotypes, and so summaries of population genetic data usually begin with this data structure. However, since the genetic variation represented in a genotype matrix was generated by mutations inherited through genealogical trees, it is often informative to think about these statistics in terms of these trees. Most population genetic summary statistics can be thought of in terms of the *allele frequencies,* i.e., what proportion of some subset of the sampled genotypes carry each allele. The frequency of any particular allele can easily be found from the location of the mutation(s) that produced it on the marginal genealogical tree, by counting the number of sampled genotypes below the mutation(s) in the tree. If many mutations have occurred at distinct sites that share a single genealogical tree, then these mutations start to give us an idea of the shape of that tree. This relationship is depicted in Figure 1, which shows the relationship between the genealogical tree and the frequencies of alleles at five different sites across a region of the genome described by that single tree. Our goal will be to generalize this sort of summary of the genotype matrix, using the underlying genealogical trees. Working with the trees serves two purposes. First, the trees are closer to the demographic processes that we are really interested in, and so allow us to reason more directly about population genetic signal and noise. Second, trees are highly efficient data structures, and allow us to compute with genetic data far more efficiently than with the genotype matrix itself.

**Figure 1:**
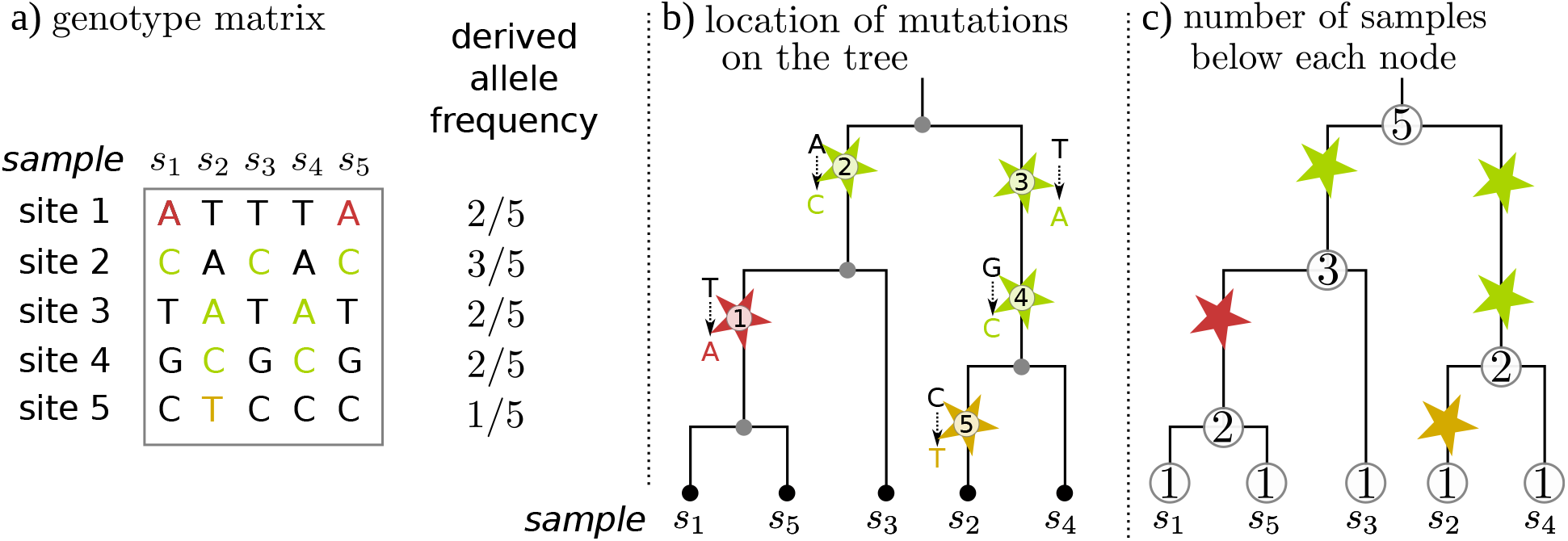
The relationship between allele frequencies and the underlying genealogical trees. **(a)** The genotype matrix, showing genotypes of five sequences (“samples”) at five separate sites, as well as the derived allele frequencies (derived alleles are shown in color, with separate colors for each genotype pattern). **(b)** The genealogical tree describing how those five samples are related to each other over this segment of genome. The samples are leaves, and each mutation is represented as a star on the genealogy, with color matching the genotype matrix, and labeled with its index in the genotype matrix and the ancestral and derived states at that site. **(c)** Each node in the tree is labeled with the number of samples at or below that node, which we think of as a *weight.* Note that this is *additive*: the weight of each node is the sum of the weights of its child nodes. Since the frequency of a derived allele is the number of samples that have inherited that mutation, the allele frequency of each mutation is equal to the weight of the node directly below the mutation, divided by the total sample size.

The operation of finding allele frequencies on a tree, depicted in Figure 1(c) is the basis of a familiar operation in population genetics: summarizing the allele frequency spectrum. Somewhat more generally, we could calculate how many of a certain set *A* of samples inherit from a particular branch in a tree by (a) assigning each of these samples weight 1 (and other samples weight zero), then (b) finding the total weight of all samples in the subtree inheriting from that branch. This gives the number of samples that would inherit any mutation falling on this branch. Suppose we want to calculate one of the linear functions of the frequency spectrum discussed by Fu [1995] and Achaz [2009], which can be described as follows. Write *p_i_*(*a*) for the frequency of allele *a* at a site *i*, and *h*(*p*) for the allele frequency spectrum — i.e., the number of alleles whose frequency is *p* in our sample — and suppose that for some function *f* (), we want to compute Σ_*p*_ *h*(*p*)*f* (*p*). Since this is a simple sum over sites, we can compute this by summing over the *L* sites in the genome and all alleles (including the ancestral allele),

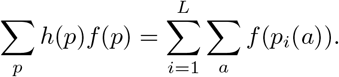

For instance, mean genetic diversity across biallelic loci is computed using this formula with *f* (*p*) = *p*(1 – *p*), omitting the usual factor of two because the sum is over *both* alleles at each site. This is because genetic diversity is defined to be the probability that two randomly chosen sequences differ at a randomly chosen site, and the probability that two randomly chosen genomes differ at a site with allele frequency *p* is *p*(1 – *p*) + (1 – *p*)*p*. If we know where each mutation has occurred on the genealogical tree at each site in the genome, we can easily compute statistics of this form by finding allele frequencies, applying the function f () to each, and summing up the results.

Our framework is somewhat more general than this, since we also define statistics that are *not* functions of the allele frequency spectrum. The small but useful generalization we make is to allow the “weights” we propagate up the tree to be numbers other than zero or one. The general procedure is depicted in Figure 2. If weights *are* all zero or one, then these weights will count numbers of samples in each part of the tree; but below we use other weights to compute the correlation between genotype and phenotype.

**Figure 2:**
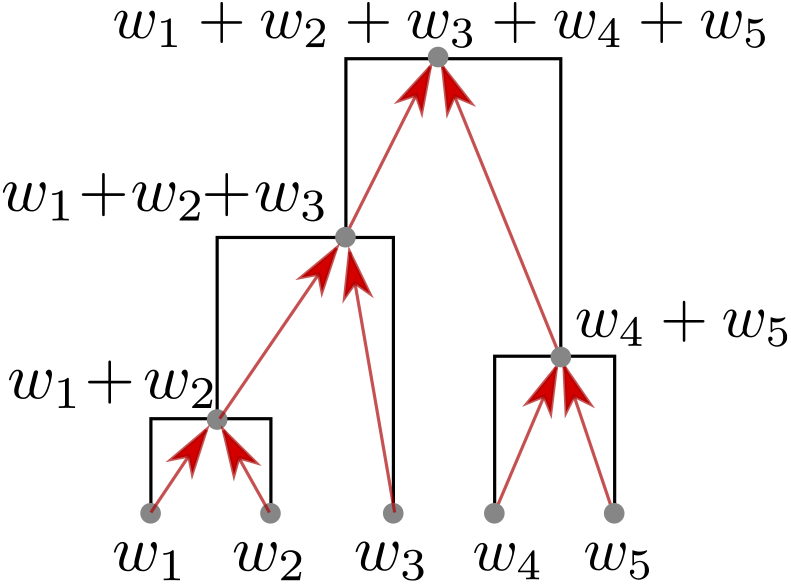
*Subtree weights* are assigned to each node by adding together the weights of any children, plus the weight of the node itself, if it is a sample. In this example, the samples are the five leaves, and have weights *w*_1_,…,*w*_5_; the total weight - also the subtree weight of the root - is *w*_total_ = *w*_1_ + … + *w*_5_. This can summarize the tree in many ways: for instance, if all weights were equal to one, then subtree weights would count the number of samples below that node. On the other hand, if *w*_1_ = 1 but other weights were zero, then subtree weights would indicate whether a node was an ancestor of sample 1. If *w_k_* was the value of a phenotype of sample *k*, then the subtree weight of a node would equal the sum of all phenotypes inheriting from that node.

Before we formally describe the statistics, we need some notation and definitions. A tree sequence describes how a set of *n* sampled chromosomes are related to each other along a (linear) genome of length *L* [Kelleher et al., 2016, 2018]. Each haplotype, modern or ancestral, is associated with a *node,* and the trees at each position along the genome have vertexes labeled by these nodes. A tree sequence describes the relationships between a special set of nodes, the *samples,* that appear in the trees at every point along the genome. The other (non-sample) nodes may not appear in every tree, as for instance if a portion of an ancestor’s genome was not inherited by any of the samples. From the tree sequence can be extracted a sequence of trees, 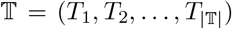 and a sequence of breakpoints 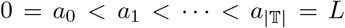, where *T_k_* describes the genealogical relationships of the samples over the segment of genome between *a*_*k*-1_ (inclusive) and *a_k_* (exclusive), and 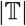 is the number of trees. We say that tree *T_k_ covers* the (half-open) segment [*a*_*k*-1_, *a_k_*), and call the length of this segment its *span,* denoted *L_k_* = *a_k_* – *a*_*k*-1_. We refer to the branches in each tree using the most recent node, so for instance a branch between an ancestor *v* and descendant *u* is associated with the node *u*. Note that although the same parent-child relationship may exist across many adjacent trees (this is called an “edge”), rearrangements of genealogical relationships due to recombination can cause the precise set of samples that inherit from that edge to differ across trees.

### Definition 1

(Sample and subtree weights). *A list of* sample weights *w assigns a numeric value w(v) to every sample node. Given these weights, the* subtree weight *X_k_* (*u*) *on tree T_k_ for a node u is the sum of all sample weights of every sample node that is descended from u in the tree (including u, if it is a sample)*:

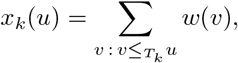

*where v <T_k_ u if u is on the path from v to root in the tree T_k_. The* total weight *is the sum of the weights over all samples: w_total_* = Σ_*v*_ *w*(*v*).

If a given node *u* is a sample and has no offspring (i.e., it is a leaf), then its “sample weight” and “subtree weight” are the same: *x_k_*(*u*) = *w*(*u*). However, these differ for any samples that are internal nodes in a tree, and so we use different notation to distinguish the two concepts. Also note that *w*_total_ is the subtree weight of the root of any tree, as long as the tree contains all the samples.

We allow vector-valued weights, i.e., *w*(*v*) may be a vector (*w*_1_(*v*),…, *w_m_*(*v*)), so that when summarizing we have access to more than one aspect of each subtree. We will often be interested in statistics defined in terms of different subsets of our samples; it is useful to have some additional notation to define these statistics. Given a subset set *S* of samples, the *indicator weights of S* are the sample weights **1**_*S*_ with **1**_*S*_(*u*) = 1 if *u* ∈ *S* and **1**_*S*_(*u*) = 0 otherwise.

### Definition 2

(Summary function). *For a set of k-dimensional weights with total weight w_tota_ι, a summary function is any real-valued function f* (*w*_1_, …, *w_k_*). *We call the summary function* strict *if f* (0) = *f* (*w_total_*) = 0.

The requirement that *f* (0) = *f* (*w*_total_) = 0 ensures that statistics do not depend on portions of the tree either not ancestral to any of the samples or ancestral to all of them. This is desirable because genetic variation within the samples cannot inform us about such parts of the tree. However, as we will see below it is sometimes useful to use non-strict summary functions.

### Node statistics

Perhaps the most natural way to summarize weights on the tree is simply to examine the values at each node. This would allow us, for instance, to count the number of samples that inherit from each node in the tree. Averaged across trees, this tells us, for each node, what proportion of the sample’s genomes were inherited from that node. Motivated by this, we define the **node statistic** for node *u* associated with summary function *f* () and sample weights *w* to be the sum of *f* () applied to the weight of the subtree inheriting from node *u* and to the remaining weight of the rest of the tree, averaged across the genome:

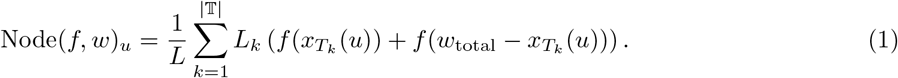

The weight *x_T_k__* (*u*) is the sum of the weights of all samples descending from node *u* in the tree *T_k_*, and *w*_total_ – *x_T_k__* (*u*) is the sum of all remaining weights. This is an average over the genome because each tree is weighted by its span, *L_k_*, and the sum is divided by the total genome length, which is the sum of all the spans: 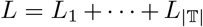. If the node statistic is computed over only a “window” of the genome then *L_k_* is replaced by the length of the portion of that window that the tree *T_k_* extends for, and *L* is replaced by the length of the window.

#### Polarization

The node statistic defined above is appropriate for statistics of all alleles at given variable site. For other statistics, one is only concerned with the derived state(s) at a site, and we define a *polarized* node statistic without the second term from our previous definition:

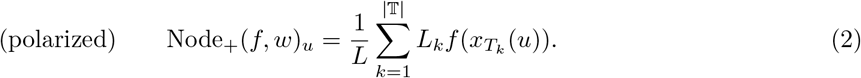

#### Example 1

(Ancestry proportions). *If w* = **1**_*S*_ *are the indicator weights of the set S of n samples, then x(u)/n is the proportion of the samples in S that inherit from u. Therefore, if f (x) = x/n, then Node*_+_(*f,w*)_*u*_ *is the proportion of the genomes of S that are inherited from (ancestor) u.*

Dividing this statistic by the total amount of the genome each node is ancestral to any samples, we would obtain the “genomic descent” statistic defined by Scheib et al. [2019].

Note that the summary function in this last example was not *strict,* since *f*(*w*_tota1_) = 0. Strictness is less important for node statistics than for the remaining classes of statistic, because Node statistics only make sense in the context of the tree sequence. However, Node statistics provide a useful bridge to the next type of statistic, which are defined directly in terms of the genotypes.

### Site statistics

Now we describe how to compute statistics from *genomes* using this framework. To do this, we assume that the genetic variation data is embedded in a tree sequence, but the trees are used only for efficiency – the results will not depend on the trees in any way. The summaries we defined above for nodes extend directly to genetic variants: just as a node weight contains information about which samples inherit from the node and which do not, so we can summarize patterns of genetic variation by summing up weights of all samples that carry a given allele. Therefore, we define *allele weights* to be the total weight of all sample nodes that have inherited that allele.

#### Definition 3

(Allele weights). *The* allele weight *for allele a at site j is the sum of the weights of all sample nodes inheriting this allele:*

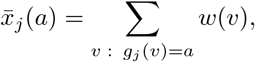

*where the sum is over all sample nodes v for which g_j_(v), the allele carried by node v at site j, is equal to a.*

If there has been only one mutation at the site, then 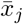 (a) is equal to the weight of the subtree inheriting from the mutation that produced *a*. In the more general case of recurrent and back mutations, 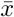 can be computed from subtree weights in a straightforward way. We then define the **site statistic** of site *j* for a summary function *f* () and sample weights *w* to be the sum of *f* () applied to the weight of every allele, *a*, found at this site:

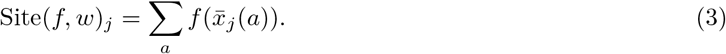

Here, we think of a “site” as a single nucleotide position on the genome. We often want to summarize statistics across regions of the genome (“windows”). To do this, we overload notation somewhat and use a subscript [*i, j*) to denote an average over the corresponding portion of the genome:

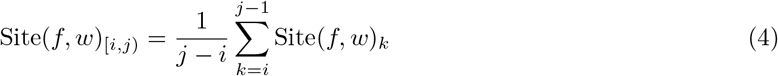

The requirement on strict summary functions that *f* (0) = *f* (*w*_total_) = 0 ensures that the sum is only affected by polymorphic sites, although we normalize by total number of sites, so that the values are comparable between different regions of the genome.

In the definition above we sum over all alleles at each site. However, sometimes it is useful to distinguish the *ancestral* allele (i.e., the allele at the root of the tree) from the remaining derived alleles. This allows statistics in principle to differentiate ancestral from derived alleles, information which is available in practice (albeit noisily), and one way to make use of this information is to sum over only derived alleles. Analogously to the above, we say a site statistic is **polarized** if we do this, defining

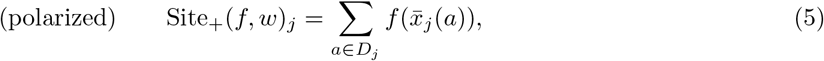

where *D_j_* denotes the set of all alleles at site *j* except the allele at the root of the tree. Note that this may not be quite what is expected, for instance, if there has been a back mutation to the ancestral allele at some point in the tree, or if there have been mutations to distinct alleles on different parts of the tree such that the ancestral allele is no longer present. However, since these situations depend on multiple mutations occurring at a single site, they are relatively rare in practice.

Having defined a general framework, we may now give concrete examples of how to compute common summary statistics.

#### Example 2

(Nucleotide diversity). *The nucleotide diversity of a group S of n samples is the average density of differences between pairs of samples, or equivalently, the average heterozygosity across positions. To calculate this statistic, let w* = **1**_*s*_, *so that x*(*u*) *gives the number of nodes in S inheriting from u, and define*

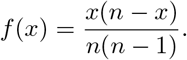

*Then Site*(*f,w*)_[*a b*)_ is mean nucleotide diversity of S in the region of the genome between a and b.

An alternative and possibly more familiar way to compute heterozygosity would be to use a polarized Site statistic with summary function *f* (*x*) = 2*x*(*n* – *x*)/ (*n*(*n* – 1)). However, this approach fails with more than two alleles per site.

#### Example 3

(Nucleotide divergence). *Now suppose we want to compute the mean pairwise density of nucleotide differences* between *two nonoverlapping groups of samples, S_1_ and S_2_, with n_1_ and n_2_ samples, respectively. As before, let w_j_* = 1_*s*_*j*__, so that *x_j_* (*u*) *gives the number of nodes in S_j_ inheriting from u, and define*

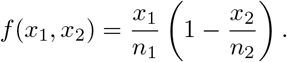

*Then Site(f, w)*_[*a,b*)_ *is mean nucleotide divergence between the two groups in the region of the genome between α and b.*

The procedure of computing divergence at a single site is shown in Figure 3; divergence over a region of the genome would sum these values across all polymorphic sites and divide by the length of the region.

**Figure 3:**
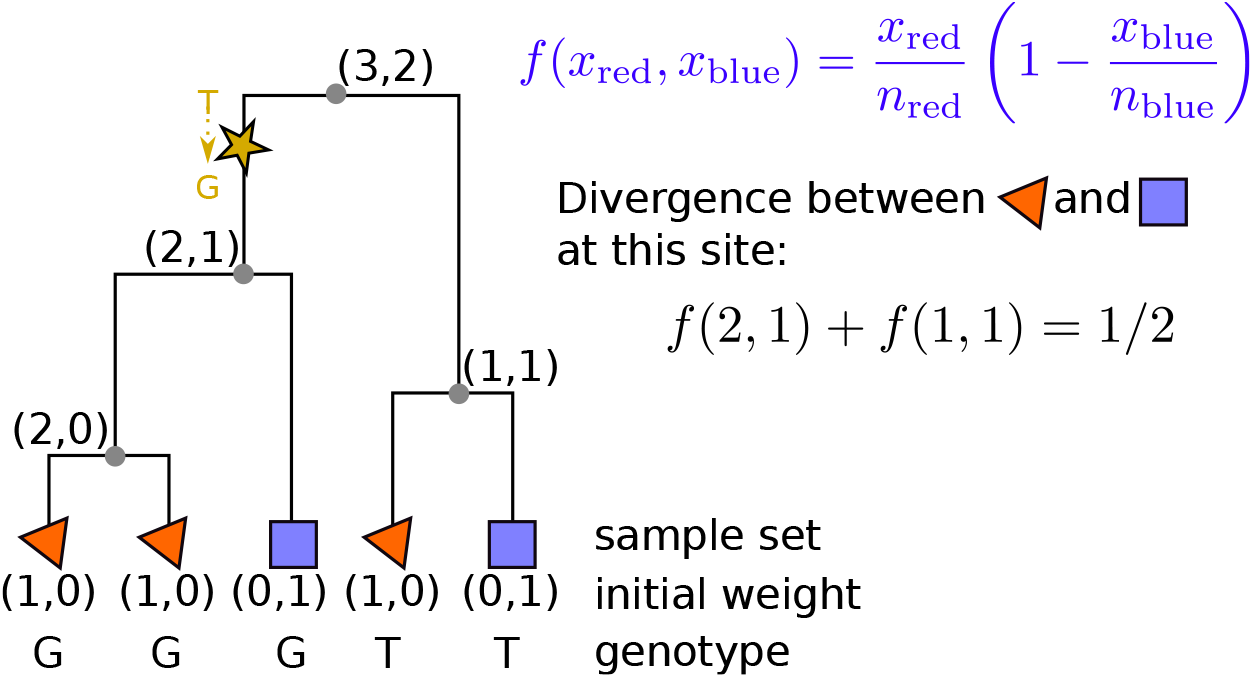
An example of computing sequence divergence between red samples (triangles) and blue samples (squares) at a single site with a T→G mutation, using the tree. Each node is labeled with subtree weights (*x*_red_, *x*_biue_), recording how many red samples (*x*_red_) and how many blue samples (*x*_biue_) lie in that subtree. The derived mutation is found in *x*_red_ = 2 out of the *n*_red_ = 3 red samples and in *x*_blue_ = 1 of the *n*_blue_ = 2 blue samples, and so the probability that a randomly chosen red node carries the derived allele but a randomly chosen blue node does not is given by the summary function, *f* (*x*_red_, *x*_blue_) = *f* (2,1) = 2/6. The complementary value, *f* (1,1) = 1/6 is the probability that the mutation separates a randomly chosen red and blue pair, but with the blue node carrying the derived allele, so the total probability that a randomly chosen pair differs at this site is *f* (2,1) + *f* (1,1) = 1/2.

#### Example 4

(Segregating sites). *Again, let w* = 1_*s*_ *for a group of n samples, and now define*

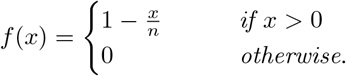

*In computing Site(f, w)*_[*a b*)_, *the argument x to the function f* (*x*) *is the number of samples in S that inherit each allele. A site with k distinct alleles that segregate in S with counts x*_1_ + … + *x_k_* = *n contributes f* (*x*_1_) + … + *f* (*x_k_*) = (1 – *x*_1_/*n*) + … + (1 – *x_k_*/*n*) = *k* – 1 *to the statistic. Therefore, Site(f, w)*_[*a,b*)_, *is the minimum number of derived mutations per unit length, which is the density of segregating sites if all sites are biallelic. (The actual number of mutations per unit length will be greater if there have been back mutations*.)

#### Polyallelic sites

The summary function in Example 4 might be surprising: the reason for its particular form is so that the statistic returns a sensible answer even when sites may have more than two alleles. There is not a consensus in the field on how to use sites with more than two alleles, but the most common practice is to reduce to biallelic data, by either discarding other sites or by marking all non-ancestral alleles as (the same) “derived” allele. In contrast, we choose to define statistics in a way that still makes formal sense for polyallelic sites, so that for instance a site with ancestral allele *A* and derived alleles *C* and *T* would have site statistic 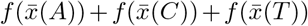. This is not the only choice, but is natural because it agrees with what is obtained by looking at pairwise haplotype differences. For instance, the definition of nucleotide divergence in Example 3 gives the probability that a randomly chosen sample from each set differ, a definition that makes sense even with polyallelic sites.

#### Example 5

(Phenotypic correlations). *Suppose that for each sample u we have a numeric phenotype, denoted z(u), and we want to compute the correlation between this phenotype and the genotype at each site. For convenience, suppose z is normalized to have mean zero and variance 1. Then, if gj is a vector of binary genotypes* (*so g_j_* (*u*) = 1 *if u carries the derived allele*), *then the covariance of z with g_j_ is just Σ_u_z*(*u*)*g_j_* (*u*) = Σ_*u: g*_*j*_(*u*)=1_ *z*(*u*), *i.e., the sum of the phenotypes of all samples carrying the derived allele, divided by* (*n* – 1), *where n is the number of samples. Since the phenotypes sum to zero, this is also equal to* – Σ_*u*: *g_j_*(*u*)=0_ *z*(*u*). *If p_j_* = Σ_*u*_ *g_j_* (*u*) *is the derived allele frequency at site j, then the variance of g_j_ is np_j_* (1 – *p_j_*)/(*n* – 1), *and so the squared correlation can be calculated as a sum across the two alleles*:

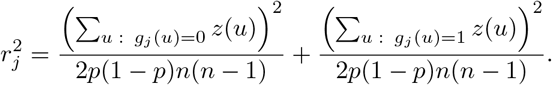

*We can compute this as a site statistic by defining w*_1_(*u*) = *z*(*u*), *and w*_2_(*u*) = 1/*n*, *and f*(*x*_1_, *x*_2_) = 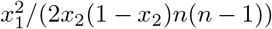: *then Site* 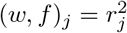 *is the squared correlation between z and the allele at site j*.

Appendix A explains how to extend the previous example to obtain the squared coefficient of genotype in the regression of phenotype against genotype with additional covariates.

#### Example 6

(Patterson’s *f*_4_). *Given four disjoint groups of samples, S*_1_, *S*_2_, *S*_3_, and *S*_4_, *Patterson’s f*_4_(*S*_1_,*S*_2_; *S*_3_,*S*_4_) *statistic [Reich et al., 2009, Patterson et al., 2012] for an allele with frequency p*_1_ *in group S_i_ is* (*p*_1_ – *p*_2_)(*p*_3_ – *p*_4_). *To rewrite this as a sum over alleles, note that p*_1_ – *p*_2_ = *p*_1_(1 – *p*_2_) – (1 – *p*_1_)*p*_2_, *and so the statistic counts with positive value alleles that split S*_1_ *and S*_3_ *from S*_2_ *and S*_4_, *and negative value ones that split S*_1_ *and S*_4_ *from S*_2_ *and S*_3_. *Therefore, if as before we let w_j_* = **1***s_i_* (*so that w_j_*(*u*) *tells us the number of samples in S_j_ descended from u*), *and write n_i_ for the number of samples in S_i_, and*

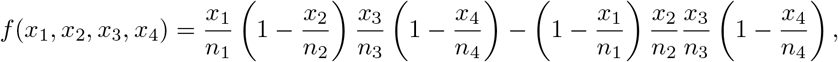

*then Site*(*w,f*)_[*a,b*)_ *is equal to Patterson’s f*_4_(*S*_1_, *S*_2_; *S*_3_, *S*_4_) *statistic for the region of the genome between a and b*.

Again, we have taken care so that this definition makes sense even with more than two alleles per site. This definition of *f*_4_ can be restated as follows: averaged across a random choice of individuals from the four groups of samples, let “BABA” be the proportion of sites at which samples 1 and 3 agree, but differ from samples 2 and 4 (which may differ from each other as well); and let “ABBA” be the proportion of sites at which 2 and 3 agree, but differ from 1 and 4; then *f*_4_ is BABA minus ABBA. In this mnemnonic, B is standing in for a *specific* allele, that must match; but A is standing in for “any allele that is not B”. This is more general than previous definitions, but agrees with them for biallelic sites.

### Branch statistics

Genetic variation is informative about many processes precisely because it tells us about the underlying patterns of genealogical relatedness. In other words, often the genomes are most useful in so far as they tell us about the trees. In the case where we actually have the trees, or a good proxy for them, it is natural to summarize them directly rather than working indirectly with the genotypes [Harris, 2019]. If we assume that no two mutations occur at the same genomic position – i.e., the *infinite sites model* – then there is a natural correspondence between summaries of genotypes and summaries of tree shape. If mutations occur at a constant rate in time and along the genome, then the *expected* number of mutations that occur somewhere along a branch of a tree over some segment of genome is equal to the mutation rate multiplied by the length of the segment and by the length of the branch. In other words, the “area” of a branch in a tree, defined as its span (right minus left endpoint) multiplied by its length (parent time minus child time) and scaled by the mutation rate, is equal to the expected number of mutations that will land on it. If the mutation rate is constant, then this gives us an isometry: tree distances measured in branch lengths versus in numbers of mutations are equal in expectation, up to a multiplicative factor of the mutation rate.

This makes it natural to define a statistic of tree shape by summing these expected contributions across its branches. We define the **branch statistic** of the *k*^th^ tree *T_k_*, obtained from summary function *f*() and sample weights *w*, to be the sum of the length of each branch multiplied by the summary function applied to its subtree weight and the remaining weight not in the subtree:

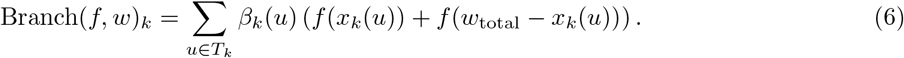

Here, *β_k_*(*u*) is the length of the branch between *u* and its immediate ancestor in tree *T_k_, w*_total_ = Σ_*u*_ *w*(*u*) is the total weight, and *x_k_*(*u*) is the total weight of the subtree of *T_k_* inheriting from *u* (as defined above). The term *w*_total_ – *x_k_*(*u*) gives the total weight *not* in the subtree of *T_k_* inheriting from *u*. The value *f* (*x*(*u*)) is the summary value that would be added to a site statistic if a single mutation occurred on the branch ancestral to *u*, and *f* (*w*_total_ – *x_k_*(*u*)) is the value that would be added due to its complementary allele, so Branch(*f, w*)_*k*_ gives the expected contribution of the tree *T_k_* to Site(*f, w*), per unit of sequence length that the tree covers. An example of this correspondence between site and branch statistics is shown in Figure 4. Here, we see how the *f*_4_ site statistic assigns a weight to each mutation depending on its frequency in each of the four sample sets, and how the corresponding branch statistic assigns the same weight to those branches.

**Figure 4:**
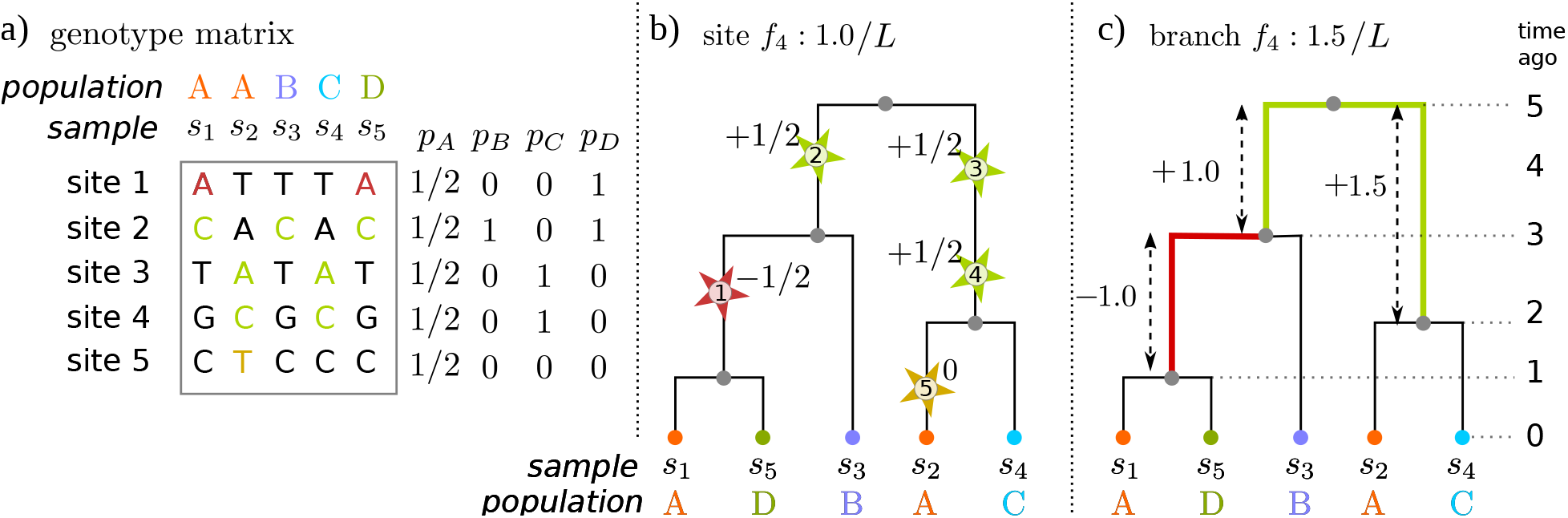
Computing the *f*_4_(*A, B; C,D*) statistic on a tree, for groups of genomes labeled *A, B, C*, and *D* (only group A has more than one sample), on the same basic example as Figure 1. **(a)** The genotype matrix, for five sampled genomes at five sites. Also shown are the derived allele frequencies in each of the four groups, denoted by *p_A_, p_B_, p_C_*, and *p_D_* respectively. **(b)** Each mutation is given weight equal to (*p_A_* – *p_B_*)(p_c_ – *p_D_*), shown as a label next to each mutation (depicted as a star), and the Site *f*_4_(*A, B; C, D*) statistic is the sum of these weights divided by *L*, the sequence length. Note that each of the mutations shown occurs at a distinct site, so there is no back mutation, and that all mutations on the same branch have the same weight. **(c)** Each branch is assigned weight equal to the weight that would be assigned to any mutation falling in it (colored lines, with colors matching the mutations of (b)), and then the branch *f*_4_ statistic is the sum over these weights, multiplied by the length of the branch, and divided by L, the sequence length. Remaining branches do not contribute, because any mutation falling on these have either *p_A_* – *p_B_* = 0 or *p_C_* – *p_D_* =0.

Equation (6) defines a statistic for a single tree, but in practice it is more useful to average branch statistics over a region of the genome, as we did for Site statistics. We do this by averaging the tree statistics over the region with probabilities proportional to the trees’ spans:

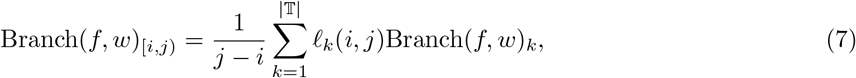

where *ℓ_k_* (*i, j*) is the length of the region in [*i, j*) that the tree *T_k_* covers (i.e., if *T_k_* covers the half-open interval [*a*_*k*-1_,*a_k_*), then *ℓ_k_*(*i,j*) = max(0, min(*j,a_k_*) – max(*i, a*_*k*-1_))). The *polarized* version of a Branch statistic is defined analogously to the Node and Site versions:

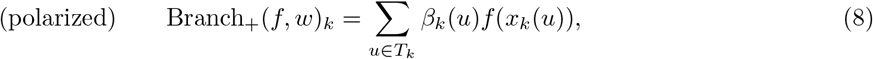

and Branch_+_(*f, w*)_[*i,j*)_ is defined using Branch_+_(*f, w*)_*k*_ as before.

#### Example 7

(Mean TMRCA). *If we take w and f exactly as in the “Nucleotide diversity” example above, then f* (*u*) *gives the probability that the branch between u and its ancestor in the tree lies on the path from one of two randomly chosen samples from S on the path up to their most recent common ancestor (MRCA). Therefore, Branch(f, w) gives the mean total distance in the tree between two samples from S, averaged across the sequence. This is twice the mean time back to the MRCA if the samples are all from the same time.*

#### Example 8

(Phenotypic correlation with pedigree). *If we take w and f as in the “Phenotypic correlations” example above, then Branch(f, w) gives the expected correlation between phenotype and any mutations appearing on the tree. This is a summary of how much local relatedness aligns with similarity in phenotype.*

The previous example might be used to leverage local relatedness to improve the resolution of association studies. Similar strategies were explored by Zöllner and Pritchard [2005] and Minichiello and Durbin [2006].

#### Example 9

(Patterson’s *f*_4_). *Suppose that the four subsets each consist of only a single sample. The summary function f* (*x*_1_, *x*_2_, *x*_3_, *x*_4_) *for the f*_4_ *statistic then assigns weight 1 to any branch that separates x*_1_ *and x*_3_ *from x*_2_ *and x*_4_, *and weight −1 to any branch that separates x*_1_ *and x*_4_ *from x*_2_ *and x*_3_. *The statistic Branch(f, w) therefore gives the difference in average lengths of these two types of branch, averaged across the genome.*

## Duality of site and branch statistics

Under a neutral, infinite-sites model of mutation with constant mutation rate across time, the expected number of mutations per branch is proportional to its length. This implies an isomorphism between “Site” and “Branch” statistics defined above, which is discussed in more detail in Ralph [2019]. For instance, the site statistic of Example 2 (genetic diversity) and the branch statistic of Example 7 (mean TMRCA) use the same summary function *f* (*x*) = *x*(*n – x*) / (*n*(*n* – 1)). These are closely related because under an infinite-sites model of mutation, two sequences differ at a site only if there has been a mutation somewhere on the branches going back to their most recent common ancestor. Therefore, if mutations occur with constant rate the expected value of genetic diversity is equal to the mutation rate multiplied by the average path distance between the two samples in the trees.

This relationship is true more generally. In fact, for any region of the genome between *i* and *j*,

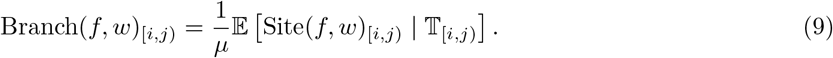

Here, 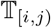 denotes the tree sequence on the interval [*i, j*), and so the expectation keeps the genealogies fixed and averages over infinite-sites mutations with rate *μ* per unit time and per unit of sequence length. Note that the expected product of two site statistics, 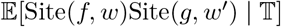 is not equal to the product of the two branch statistics, *μ*^2^Branch(*f, w*)Branch(*g, w*′), because they are not independent. However, it is always possible to define a branch statistic that gives the expected value of the product, as described in Ralph [2019].

A Site statistic can therefore be thought of as an estimator of its corresponding Branch statistic, which is itself a summary of local tree shapes. Because we have normalized these statistics by the length of the genome under consideration, both sides of this equation are in units of *time*: Branch statistics give mean weighted branch lengths; Site statistics give mean densities of mutations per unit of sequence length, which divided by the mutation rate *μ* is converted to time. Let’s unpack the assumptions here: what exactly is the mutational model? First, we are taking the mutation rate to be *μ*, i.e., the expected number of mutations that occur on a region of the genome of length *ℓ* over *t* units of time is equal to *μℓt*. Second, we are assuming that the probability of per-site mutation is low enough that no two mutations occur at the same site – the fact that they do, occasionally, means that this is an approximation. Third, we are assuming that mutation rates are constant through time and across the genome. Of course, the statement remains true if we can measure distance along the genome and time in a way that mutation rates are constant, but how these vary is generally unknown.

In this view, site statistics are noisy approximations to the corresponding branch statistic – but, how noisy? How big is the contribution of mutation to the overall sampling variance of a statistic? The law of total variance partitions the variance of a site statistic into the contributions of noise from demography and mutation:

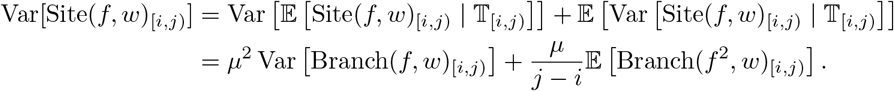

The first term is the variance of the expected Site statistic given the trees, which by duality is the variance of the Branch statistic, i.e., the contribution of demographic noise, including genetic drift. The second term, the expected variance of the statistic given the trees, is the contribution to variance of mutation, and can itself be written as a Branch statistic, using the same weights and the summary function *f* (*x*)^2^ [Lemma 2 in Ralph, 2019]. The reason for this is straightforward – the Site statistic is a sum across branches of the number of mutations on the branch multiplied by a summary, *f* (*x*). Given the trees, these numbers of mutations are independent and Poisson, and the variance of the product of the constant *f* (*x*) and a Poisson random variable with mean *m* is *mf* (*x*)^2^. The sum is divided by the length of the region, *j* – *i*, and so the variance is divided by (*j – i*)^2^, but one factor of (*j* – *i*) is absorbed by the definition of the branch statistic.

We examined the contributions of mutation and demography to noise in two simulations where population genetic (Site) statistics are important in making inferences: detecting recent selective sweeps, and detecting introgression. Figure 5 shows windowed diversity along the chromosome following a few selective sweeps. The top two plots compare Branch diversity – i.e., as computed only with tree shape – to Site diversity computed from sequences generated by 20 independent assignments of mutations to the same tree sequence, with mutation rates 10^-9^ and 10^-8^, respectively. We see that as the mutation rate increases, the signal of decrease in diversity around swept loci becomes more clear, and Site diversity approaches Branch diversity. These were computed using the entire population of 1,000 individuals; how does sampling variance contribute? Not much, it turns out – the bottom plot shows both Site and Branch diversity computed from 20 nonoverlapping groups of 100 samples. Neither Site or Branch diversity vary much between these samples, implying that the subsample gives us a good estimate of the whole-population values of each. However, as we see in the top figure, whole-population Site diversity is itself only a quite noisy estimator of Branch diversity. (Note, however, that statistics other than diversity may vary much more between subsamples.) The same things are shown in Supplementary Figure S1 for a simulation with 10,000 individuals, which shows similar patterns. Only the last of these plots is possible to directly observe in real data: in the top two plots, the spread of independent replicates of mutational noise (black lines) about their expectation based on the tree sequence (red line) is unobservable, although estimable.

**Figure 5:**
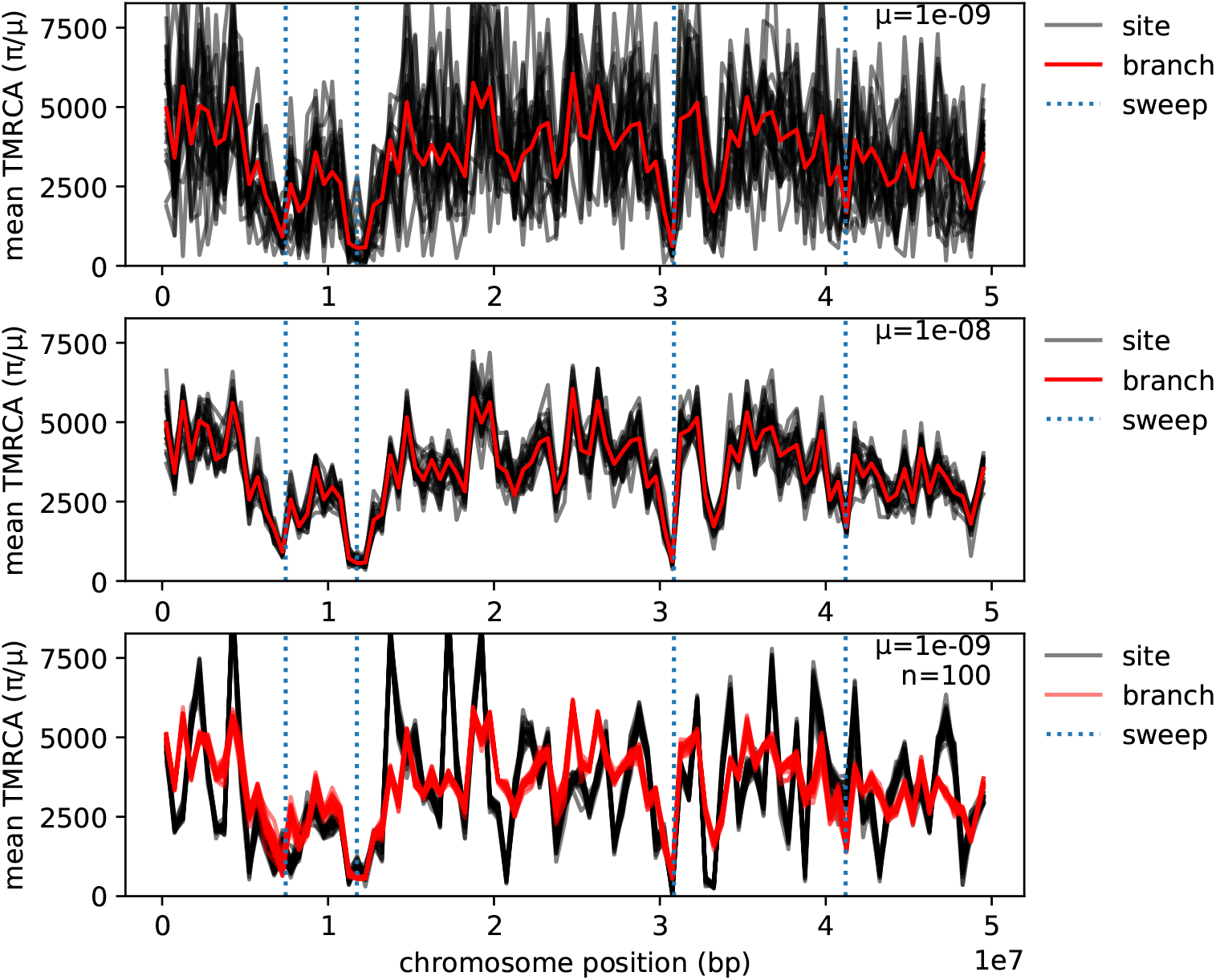
Mean genetic diversity and time to most recent common ancestor in 500Kb windows along a 50Mb genome (0.5 Morgans) following several selective sweeps. In each case, “site” is mean genetic diversity (Tajima’s *π*) divided by mutation rate, and “branch” is the corresponding Branch statistic. The tree sequence was produced by simulating mutations under positive selection with mutation rate 10^-12^ in a population of size 1000 for 400 generations using SLiM, followed by recapitation with *N_e_* = 1000 [Haller et al., 2018]. The selected alleles at the marked sites have selection coefficients between 0.08 and 0.25, and are at frequencies 96.8%, 100%, 100%, and 82.6% in the final generation, respectively. All curves use the same tree sequence, including selected mutations, but with additional neutral mutations added. **(top)** Diversity within the entire population, computed as a Site statistic from 20 independent assignments of mutations to the same tree sequence with mutation rate *μ* = 10^-9^. **(middle)** As in the top panel, but with mutation rate *μ* = 10^-8^, showing that as mutation rate increases, the Site statistic (divided by μ) converges to the Branch statistic. **(bottom)** Site and Branch diversity within 20 disjoint samples of size 100 each from a *single* assignment of mutations to the tree sequence with mutation rate μ = 10^-9^.

As another example, we simulated an “admixture” scenario: a first population of size *N* = 1000 splits into two of equal size, then after *N* generations, the second population splits, after another *N* generations, the third population splits again, and for the final *N* generations populations 2 and 3 have per-capita migration rates of 4/*N* to each other. We expect a significantly negative f_4_(1, 2; 3,4) in this situation, which we indeed find (the genome-wide mean Branch *f*_4_ is around −700 generations, as is the Site *f*_4_ divided by mutation rate; see Supplemental Figure S2 for a plot along the genome). Using various sample sizes, mutation rates, and window sizes, we then calculated this *f*_4_ statistic in windows along a 100Mb genome, and show the standard deviation of both Site and Branch statistics across windows in Figure 6. Since the genome is uniform (no selection or variation in recombination or mutation), this standard deviation is a measure of noisiness. As expected, Site statistics are noisier than Branch statistics, by a factor that depends mostly on the ratio of mutation to recombination rates. The results suggest that Branch statistics would provide substantially better resolution at small scales along the genome, especially if mutation rate is lower than recombination rate. However, in practice imperfect estimation of the tree sequence would introduce additional noise, so it remains to be seen if the improvement could be made in practice.

**Figure 6:**
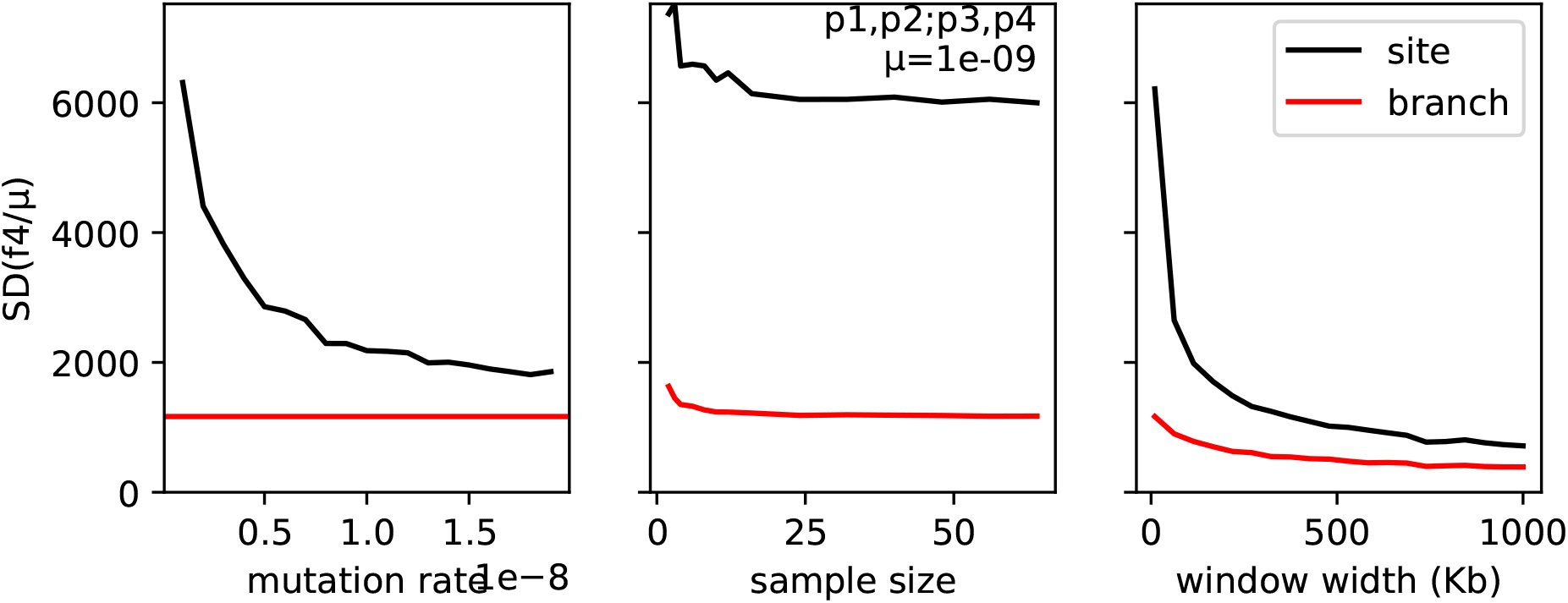
Standard deviation of Site (black) and Branch (red) *f*_4_(1, 2; 3,4) in windows along a 100Mb genome, as a function of **(left)** mutation rate, **(middle)** sample size, and **(right)** window width. The recombination rate was 10^-8^, and unless otherwise stated, the mutation rate was 10^-9^, the window width was 10Kb, and the entire population (1000 diploids in each of four populations) was used.

## Application to 1000 Genomes tree sequences

We do not know the true genealogies underlying real data, but recent methods are available to estimate them at scale [Kelleher et al., 2019, Speidel et al., 2019]. In Figure 5, we showed that Branch and Site statistics matched well in simulated data. However, these simulations make many simplifying assumptions, and, moreover, the underlying tree topologies and branch lengths are exactly correct (which inference methods can only hope to approximate). To evaluate the correspondence between Site and Branch statistics in trees inferred from real data, we calculated statistics for chromosome 20 of the 1000 Genomes dataset [1000 Genomes Project Consortium, 2015] using dated trees estimated by Relate [Speidel et al., 2019]. (Although Relate estimates a succession of essentially independent marginal trees rather than a succinct tree sequence, the output can be converted to a tree sequence, and since the statistics considered here are “single site” the distinction is not important.) Specifically, we calculated diversity using the Site statistic described in Example 2, and compared this to the dual Branch statistic (Example 7) in 1Mb windows in each of the five continental groupings. All calculations were done only using the portions of the chromosome passing the 1000 Genomes Phase 1 “strict” mask for sequencing accessibility. As shown in Figure 7, the ratio of Site and Branch diversity is relatively constant, hovering around 2.5 × 10^-8^. This is somewhat higher than typical estimates for the average genome-wide human mutation rate [e.g., 1.45 × 10^-8^, Narasimhan et al., 2017], which suggests a mis-calibration in inferred tree times. These trees were inferred using Relate with the *N_e_* fixed to 15,000; if trees are instead jointly inferred in Relate along with an *N_e_*, with mutation rate fixed, then indeed the ratio of Site to Branch diversity averages to the (externally specified) mutation rate (Figure S3). On the one hand, some amount of constancy of this ratio is expected, since Relate estimates branch lengths in part assuming that mutations fall at a constant rate through time. (But, note that Speidel et al. also showed signal of changing mutation rates, pointing the way towards a more general method.) On the other hand, since estimated node ages are shared across long regions of the genome, a tight agreement may not be possible with poorly inferred trees. So, the relative constancy of the ratio of Site to Branch diversity suggests that Relate is doing a good job at inferring trees, but what to make of the twofold variation in this ratio? Answering this question requires a deeper understanding of the processes that shape genetic diversity along the genome.

**Figure 7:**
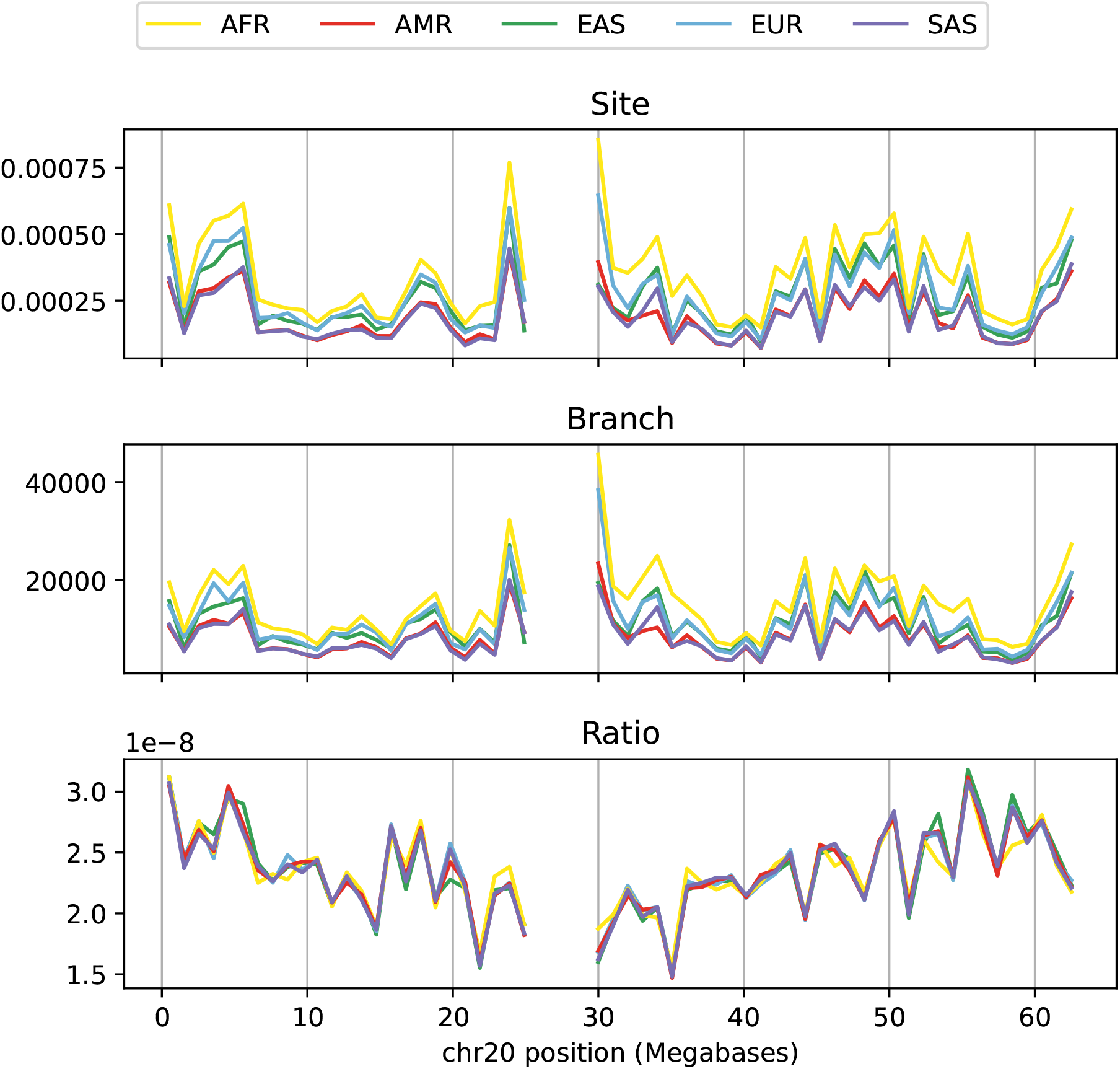
**(top)** Mean sequence diversity in 1Mb windows along human chromosome 20 of the 1000 Genomes Project, in units of mean number of pairwise differences per nucleotide. **(middle)** The dual Branch statistic as calculated from trees inferred by Relate [Speidel et al., 2019], in units of generations. **(bottom)** The ratio of the Site statistic to the Branch statistic. Separate curves are plotted for continental groupings: AFR, Africa; AMR, America; EAS, east Asia; EUR, Europe; SAS, south Asia.

In practice, we do not expect Branch and Site diversity to agree exactly, because of local differences in mutation rate and intensities of selected mutations. For instance, regions in which a high proportion of potential mutations are deleterious are expected to have a lower diversity for two reasons: first, the deleterious mutations themselves are less likely to be found, and second, the indirect effect of deleterious mutations on nearby sites reduces typical tree height and thus diversity at even neutral, linked sites [Hudson, 1994, Charlesworth et al., 1997]. The first effect would cause Site and Branch statistics to differ, because it effectively reduces the mutation rate in the region and violates the assumption of independence of mutations given the trees. However, the second effect does *not* affect the correspondence between Site and Branch statistics, since it is mediated by tree shape. It is generally unknown how much mutation rate varies along the genome [but see Supek and Lehner, 2015] or how dense targets of selection are [Leffler et al., 2012]. For this reason, it is very interesting to see how close a correspondence between Site and Branch statistics it is *possible* to obtain – it is tempting to say that the best tree sequence would obtain as tight a match between Site and Branch statistics as possible, with remaining variation explained by direct selection and/or mutation rate variation.

A final crucial assumption underlying the duality between Site and Branch statistics is that mutations can be treated as independent of the trees themselves. This is clearly not always true – for instance, a sweeping beneficial mutation strongly affects the local trees (as in Figure 5) – but seems likely to be approximately true. Ralph [2019] contains more discussion of variation in mutation rate, but more work needs to be done, particularly on models with large proportions of sites under selection.

## Algorithm and implementation

In this section we describe and analyze the algorithm used to compute Branch statistics. This is a generalization of the algorithm used to maintain the number of samples from a given set that derive from each node in a tree sequence [Kelleher et al., 2016, Algorithm L]. The central problem shared across Site, Branch and Node statistics is to efficiently maintain the “state” *x*(*u*) (where *x*(*u*) is the sum of the weights for all samples descending from, and including, node *u*; see Figure 2). As we move along the sequence, the trees change when branches are inserted and removed. By carefully ordering these insertions and removals, we ensure that this state can be correctly maintained using only small adjustments for each tree. Computing the statistics is then a straightforward matter of applying the summary function *f* to the node states and aggregating the results appropriately for the particular statistics mode and windowing options.

Formally, we represent a tree sequence using a set of tables [Kelleher et al., 2018]. Each node describes a particular haplotype (i.e., one of the two genomes of a diploid individual), and details about these haplotypes are stored in a node table, 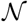. The row 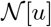 contains all information about the node *u*, and rows are indexed from zero such that 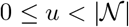. In particular, the birth time for *u* is given by 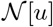.`time`. Information about how nodes relate to each other along the genome is encoded in an edge table, 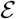. For a given row 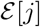.`child` and 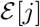.`parent` define a parent-child relationship between two nodes in a set of contiguous trees along the genome. 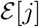.`left` then denotes the left-most (inclusive) and 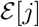.`right` the right-most (exclusive) genome coordinates over which this branch exists. The order in which edges are inserted and removed is determined by the “index vectors” **i** and **o** [Kelleher et al., 2016]. The “edge insertion” vector **i** gives the ordering of edges sorted by left endpoint, and among edges with the same left endpoint, sorted so that edges closer to the root appear later. The “edge removal” vector **o** is similar, except gives the ordering of edges by right endpoint and with edges closer to the root appearing sooner. As we move along the tree sequence, the topology of the current tree is recorded in the “parent vector” *π*: the parent of each node *u* is the node *π_u_*, with *π_u_* = – 1 if *u* is a root. Similarly, we maintain the branch length vector *β* by setting 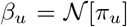.`time` 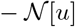.`time` and *β_u_* = 0 if *u* is a root. As evaluating the summary function *f* is an expensive operation, we also maintain the vector *F* such that *F_u_* = *f* (*x_u_*). For notational simplicity here we assume that weights and summary functions are scalars, but the extension to vectors is trivial. We also assume that we are computing the statistic across the full span of the tree sequence; the extension to multiple windows is routine. See below for a description of the steps in words, and Figure 8 for an example of the internal state of the algorithm.

### Algorithm B

(*General Branch statistics*). Given a list of *n* sample nodes *S*, corresponding weights *w* and summary function *f*: ℝ → ℝ, compute the span-normalized statistic *σ* over a tree sequence with sequence length *L* defined by the node and edge tables 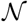 and 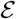. We assume that the index vectors **i** and **o** have been precomputed.

**B1.** [Initialization.] For 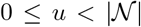 set *β_u_* ← *x_u_* ← *F_u_* ← 0 and *π_u_* ← –1. Then, for 0 ≤ *j* < *n* set *u* ← *S_j_*, *x_u_* ← *w_j_* and *F_u_* ← *f* (*x_j_*). Finally, set *s* ← *σ* ← *j* ← *k* ← *t_l_* ← 0.
**B2.** [Terminate.] If 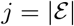 return *σ/L* and terminate.
**B3.** [Edge removal loop.] If 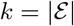 or 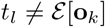.`right` go to B6.
**B4.** [Remove edge.] Set u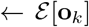.`child`, 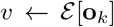.`parent` and *k* ← *k* + 1. Then set *s* ← *s* – *β_u_F_u_, π_u_* ← − 1 and *β_u_* ← 0.
**B5.** [Propagate loss.] While *v* ≠ – 1, set *s* ← *s* – *β_v_F_v_, x_v_* ← *x_v_* – *x_u_, F_v_* ← *f* (*x_v_*), *s* ← *s* + *β_v_F_v_* and *v* ← *π_v_*. Afterwards, go to B3.
**B6.** [Edge insertion loop.] If 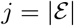 or 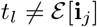.`left` go to B9.
**B7.** [Insert edge.] Set 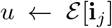.`child`, 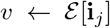.`parent` and *j* ← *j* + 1. Then set 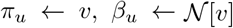.`time` – 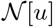.`time` and *s* ← *s* + *β_u_F_u_*.
**B8.** [Propagate gain.] While *v* = – 1, set *s* ← *s* – *β_v_F_v_, x_v_* ← *x_v_* + *x_u_, F_v_* ← *f*(*x_v_*), *s* ← *s* + *β_v_F_v_* and *v* ← *π_v_*. Afterwards, go to B6.
**B9.** [Tree loop tail.] Set *t_r_* ← *L*. If 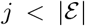 set 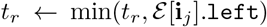. Then, if 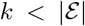 set 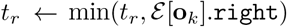 Finally, set *σ* ← *σ* + (*t_r_* – *t_l_*)*s, t_l_* ← *t_r_* and go to B2.

Algorithm B (named “B” for “Branch”) begins by setting the initial state for *π*, *β*, *x* and *F* for each node in the tree sequence, and then sets the values of *x_u_* and *F_u_* for each of the samples *u* in *S*. (The initial state is the “empty forest”, where no nodes are connected to any others.) Afterwards, we set our “running sum” *s* and output statistic *σ* to zero, along with the “tree left” variable *t_l_*. The *j* and *k* variables are used to keep track of our position in the edge insertion and edge removal indexes, respectively. After completing initialization in B1, we enter the main tree loop in B2 which is run once for each tree in the sequence. As we are processing each tree, we keep track of the edges that need to be processed with the *t_l_* variable, which stores the left-most genome coordinate in the current tree. The first thing we do for a new tree is to remove any edges that were in the previous tree and are not in the current tree. These must be the edges in which the right coordinate is equal to *t_l_*, and so B3 loops over these edges using the edge removal index **o**. Step B4 then removes the branch *u* ↦ *v* corresponding to a single edge, by subtracting its contribution to the running sum *s*, setting the parent of *u* to –1 and its branch length to 0. As shown in Figure 8B, removing an edge will result in the subtree rooted at *u* becoming disconnected from the rest of the tree. Step B5 ensures that the state of the rest of the tree is correctly maintained by propagating the loss of the state *x_u_* up the tree from *v*. Thus, for each node *v* that was a direct ancestor of *u* in the previous tree, we first subtract its contribution from the running sum s and remove the contribution of *x_u_* from its state. Since the state of *v* has changed, we must recompute the value of our summary cache *F_v_*, and then finally add the new contribution of *v* to *s*. (For example, note the changes in *x* and *F* for nodes 7 and 8 in Figure 8B.)

**Figure 8:**
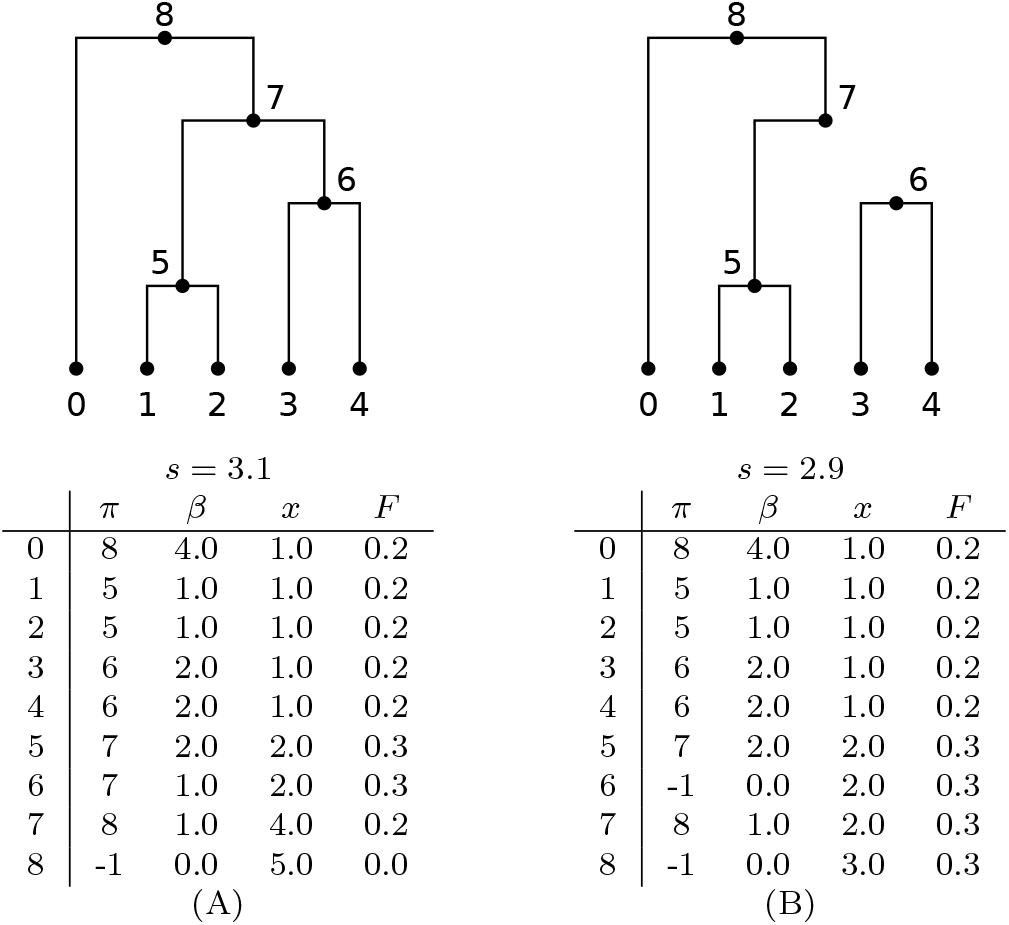
Illustration of the internal state for algorithm B for *w* = **1**_*S*_ and *f* (*x*) = *x*(5 – *x*)/20 as in Example 2. (A) Immediately before we remove the edge joining 6 to 7 and (B) immediately after. See the text for descriptions of the arrays encoding the state. Also shown is the value of the running sum, *s* = Σ_*u*_ *β_u_F_u_*.

Once all of the edges with right coordinate equal to *t_l_* have been removed in steps B3-B5, we then insert edges that start in the current tree (i.e., with left coordinate *t_l_*) in steps B6-B8. We update the state to account for the new branch *u* ↦ *v* by setting the parent of *u* to *v*, computing the branch length of *u* and adding the new contribution of *u* to *s* in B7. Then, as adding a new edge can connect the subtree rooted at *u* to the larger tree at *v*, we propagate the gain of *x_u_* up the tree in step B8 (note the symmetry with step B5). Finally, once we have removed and inserted all of the relevant edges, we are ready to add the contribution of the current tree to the overall statistic, *σ* and move on to the next tree in step B9. To do this, we first compute the right-hand coordinate of the tree *t_r_* (a process slightly complicated by the possible presence of “gaps” in the tree sequence, spanned by no edges). Then, we add the running sum s weighted by the span of the current tree *t_r_* – *t_l_* to *σ*, and return to B2 to process the next tree. If we have reached the end of the sequence, we then divide *σ* by the sequence length to normalize and exit.

To analyze Algorithm B we will assume that the tree sequence has been simulated under the standard coalescent with a sample of size *n* and population scaled recombination rate *ρ* = 4*N_e_r*, where *r* is the mean number of recombinations per chromosome per unit of time. Under these conditions, there are *O*(*n* + *ρ* log *n*) edges in the tree sequence [Kelleher et al., 2016]. Clearly, each edge is examined exactly once in steps B6 and B7 in order to create the trees, and at most once in steps B3 and B4 (we do not remove the edges for the final tree). Additionally, each edge that we insert or remove after the initial building of the tree may incur the cost of traversing up the tree as far as the root in steps B5 and B8. As coalescent genealogies are asymptotically balanced [Li and Wiehe, 2013], the expected number of nodes on the path to root is log_2_ *n*. Therefore, the overall running time of Algorithms B is of order *n* + *ρ* log(*n*) log_2_(*n*), which is

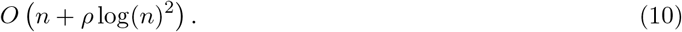

Computing a Site statistic will have the same complexity, as the same internal state is maintained, and the number of segregating sites is proportional to the number of edges (with constant equal to the ratio of mutation to recombination rate). The asymptotic analysis here assumes that we traverse upwards to root for each edge, but this is not the case. Edges are removed in oldest-first order, guaranteeing that the deepest node any subtree being modified is processed first. Therefore, this subtree will be disconnected from the main tree, ensuring that we traverse upwards all the way to root only once. Similarly, edges are inserted in youngest-first order, ensuring that subtrees are only reconnected to the main tree when they are fully built.

The tree sequence describes trees that change as one moves along the genome, and so is a special case of a *dynamic graph,* also called a *graph timeline* [Lacki and Sankowski, 2013]. Much of the work on dynamic graphs is focused on connectivity, e.g., maintaining a minimum spanning tree [Eppstein, 1994, Eppstein et al., 1997, Holm et al., 2001], but development of parallel algorithms for more general operations on large dynamic graphs is an active area of research [e.g., Srinivasan et al., 2018]. An interesting direction for future work is to develop parallel versions of tree sequence algorithms such as Algorithm B.

## Efficiency

We used coalescent simulations to compare the performance of calculating Tajima’s [1989] *D* statistic from tree sequences to calculating it from integer matrices containing genotypes at all variable positions. Figure 9 shows that the tree sequence methods implemented in `tskit` processes variant data substantially faster than matrix-based approaches once the sample size is above *n* ≈ 500 haploids – three times faster for one thousand haplotypes, growing to nearly three orders of magnitude faster for one million haplotypes.

**Figure 9:**
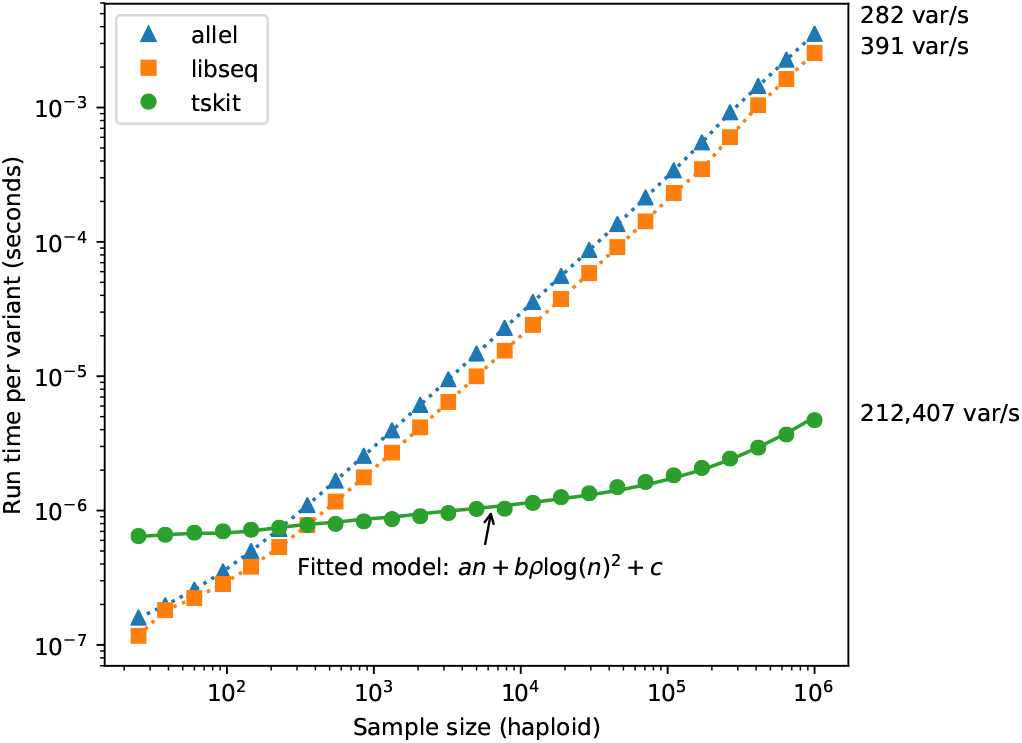
Comparison of time required to compute Site statistics between matrix and tree based methods. For each sample size, a single replicate was obtained using `msprime` with scaled neutral mutation rate *θ* = 4*N_e_Lμ* and scaled recombination rate *ρ* = 4*N_e_r* both equal to 10^4^ and *N_e_* = 10^4^, where *Lμ* and *ρ* are the total mutation and recombination rate across the simulated region per generation, respectively. For each replicate, Tajima’s *D* [Tajima, 1989] statistic was calculated using `tskit, libsequence` [Thornton, 2003], and `scikit-allel` [Miles and Harding, 2017]. The solid green line shows the result of fitting the timing data to a model based on the expected complexity of the algorithm used by `tskit` (Equation 10).

The expected number of mutations for each replicate is 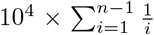, and thus ranges from ≈ 32, 000 to ≈ 138, 000 [Watterson, 1975]. Figure 9 only considers the time required to calculate the statistic once the data is present in each library’s native format. For the largest samples size of *n* = 10^6^, the matrix is approximately 200 gigabytes, and thus not practical on many systems. The largest tree sequence simulated required less than one gigabyte of memory.

To determine how well our theoretical model of time complexity predicts the running time in `tskit`’s implementation of the Site statistics algorithm, we fit a model based on Equation (10). Figure 9 shows that theoretical predictions match the observed running time very well. As the overall complexity is *O*(*n* + *ρ*log(*n*)^2^), for sufficiently large *n* the initial term (representing the cost of building the first tree) will come to dominate. In our simulations, we can see this happening at around *n* = 10^5^, where there is a noticeable uptick in the time required per variant. For longer genomes (i.e., larger values of *ρ*), this cost is amortized over more trees and is less apparent.

It should be noted here that this example based on simulated data represents a best-case scenario in terms of the performance advantages of `tskit` over the matrix-based methods. For real data, inferred tree sequences currently contain substantially more edges than we would expect based on simple neutral theory [Kelleher et al., 2019], and therefore computing statistics will not be as efficient as for an equivalent simulation. Tree sequence inference methods are in their infancy, however, and it is likely that as they improve the number of edges required to encode data will be reduced. For data simulated natively in tree sequence format via packages such as `msprime` [Kelleher et al., 2016], `SLiM` [Haller et al., 2018, Haller and Messer, 2019] and `fwdpy11` [Thornton, 2014], however, the advice is straightforward. Computing statistics using the algorithms in `tskit` will always be more efficient than decoding the genotype matrix, importing it into another package and computing from the matrix.

## Data and code accessibility

All methods described here are implemented in Python and C in the package `tskit`, available from https://github.com/tskit-dev/tskit under the terms of the MIT license. All code used to produce the figures in this paper is available from https://github.com/petrelharp/treestats_ms.

## Discussion

In this paper we have described a general framework for summarizing genetic variation and the underlying genealogies that encompasses many standard population genetic statistics. Many of these statistics are functions of the joint allele frequency spectrum, but the framework is more general and can be used, for example, to quantify associations between genotypes and phenotypes. This generality greatly reduces the software development effort in implementing statistics efficiently, and it also allows users to easily explore new classes of statistics. The range of statistics available to describe variation in a single exchangeable sample (e.g., an isolated population) are well-understood [Achaz, 2009, Ferretti et al., 2017], but there are much larger and less well-explored classes of statistics describing genetic variation between many populations or across geographical space. The statistics defined here are all additive along the genome: if we have computed a statistic in equal-size windows along a chromosome, then the value of the statistic for the entire chromosome is equal to the average of the values in those windows. Some commonly-used statistics (e.g., *F_ST_* or Tajima’s D) are not additive, but are ratios of additive statistics, so can be easily computed in this way. Extensions to statistics involving the pairwise joint distribution of genotypes across sites [Hudson, 2001] such as linkage disequilibrium are planned for future work. Haplotype-based statistics may require different classes of algorithms.

The most obvious application of these methods on current practice is to improve efficiency of existing pipelines, as tree sequences allow storage and processing of large genomic data sets with orders of magnitude less space and time than standard matrix-based methods. All statistics described here (and more) are implemented in the rigorously tested `tskit` library, which provides a suite of tools for working with tree sequences in Python and C. Large-scale simulations are useful in many contexts [e.g., Martin et al., 2017, Browning et al., 2018, Galloway et al., 2020] and the ability to quickly compute a wide range of statistics from these (previously prohibitively large) datasets opens up new possibilities. At smaller scales the statistics are still highly efficient, and avoiding the cost of exporting simulated data to a genotype matrix will in practice greatly speed up inference based on summaries computed from simulated data [Beaumont et al., 2002, Csilléry et al., 2010, Schrider and Kern, 2018]. Efficient simulations coupled with the framework developed here allow us to explore the full distribution of summaries along the genome, which has important applications in (e.g.) speciation genomics [Lohse, 2017]. The ability to efficiently compute statistics from real data is, of course, also welcome.

The correspondence between genome sequence and properties of the underlying genealogies we have used here is well-known, and is the basis for several inference methods [Becquet and Przeworski, 2007, Beeravolu et al., 2018]. The general framework that we have defined, however, codifies this relationship in a mathematically elegant and computationally efficient way, and may lead to new perspectives on well-studied problems. One way to use the duality between Site and Branch statistics is to answer: what aspect of tree shape is a particular population genetics statistic summarizing? This can help when interpreting results, especially if reality may not fit an idealized model of separate populations. However, methods to infer tree sequences from real data make it now possible to work in the other direction: instead of calculating (Site) statistics from the genotype matrix, we can calculate precisely analogous (Branch) statistics from the trees themselves, thus hopefully bypassing the extra layer of noise induced by mutation. How well this works will depend on the ability of inference methods to estimate the true tree sequence. This might plausibly produce less noisy estimates because tree sequence inference should use the signal from nearby patterns of variation to interpret relationships at a given site, thus transforming the simple binary split induced by a SNP into a much richer source of information. Furthermore, if tree sequence inference can be made insensitive to local variation in mutation rate, calculation of Branch statistics would provide a summary method that is not affected by this potentially important confounding factor. Similarly, if tree sequences inferred from genotype array data [Kelleher et al., 2019] are unbiased, then Branch statistics could provide a way to estimate genome-wide quantities without ascertainment bias. This procedure would be similar to imputation, and would likely face similar challenges.

Genomic data are naturally distributed on two axes: along the genome, and across geography. The tree sequence extends this to a third axis: time. A great many methods in population genetics aim to describe aspects of history, and accurate (or at least unbiased) tree sequence inference may shift the focus of these problems from inference to descriptive analysis. The methods developed here distribute the contribution to various statistics across each tree, and so could also be used to summarize contributions to various statistics across time. This could provide, for instance, the time distribution of mutations or branches contributing positively or negatively to f4 statistics of introgression, enabling historical interpretations of these signals. The computational toolbox of population genetics is still mostly composed of statistics originally designed for analysis of a handful of loci in a small number of discrete, mostly separated populations. Both our data and our understanding of the world have moved beyond this. We hope that the tools developed here will help make it possible to visualize and analyze genetic variation and genealogical relatedness along the genome, across space, and through time.

## Acknowledgments

Thanks to Georgia Tsambos, Graham Coop, Andrew Kern, Nick Patterson, Kelley Harris, Aaron Ragsdale, Konrad Lohse, and Wilder Wohns for useful suggestions and to Leo Speidel for providing the inferred 1000 Genomes trees. This work was supported by NSF grant #1262645 (DBI) to PLR and by NIH grant R01-GM115564 to KRT. JK was supported by the Robertson Foundation and Wellcome Trust grant 100956/Z/13/Z to Gil McVean.

## Author contributions

Designed the study: PR, JK. Wrote the code: PR, JK. Carried out the experiments: PR, KT, JK. Wrote the paper: PR, KT, JK.

## A Linear regression

Let *h* be a trait, *Z* be a matrix of covariates, and *g* be a vector denoting inheritance (so, with *n* samples, *h* and *g* are both *n*-vectors and *Z* is an *n* × *k* matrix). We would like to find the coefficient of *g* in the linear regression of *h* against *g* and *Z*, without doing full multivariate regression for every new *g*, using the fact that *Z* is always the same. Suppose that *Z^T^Z* = *I* and that the vector of all ones is in the span of the columns of *Z*, although in the implementation we post-process *Z* to make this the case. Then, let a be the number and *b* be the *k*-vector satisfying

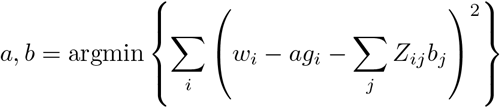

Our goal is to compute *a*. Writing this in block matrix notation, *a* and *b* minimize

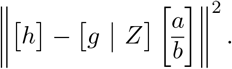

Letting *B* = [*g*|*Z*], the solution to this is

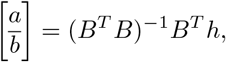

as long as *B^T^B* is invertible (which we assume to be the case). Let *m* = *g^T^g* be the number of alleles in the sample coded 1, let *u* = *g^T^Z* be the vector giving sums of the covariates of all samples carrying the allele, and

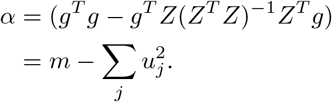

*α* is the norm of the component of *g* not in the subspace spanned by the columns of *Z*, so if *α* = 0 then we want to return *a* = 0.

Otherwise, by the inversion formula for a block two-by-two matrix, since we have assumed that that *Z^T^ Z = I*,

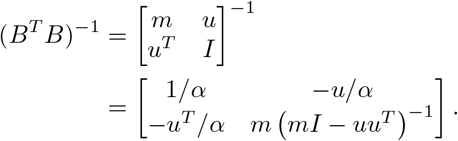

Now, the regression coefficient we seek is, with *h_g_* = *g^T^ h*,

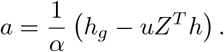

To compute this in the framework above, first add a column of 1s to the covariates *Z*, then decorrelate the resulting matrix, so that now *Z^T^Z* = *I*. Then, put this normalized version of *Z* into the first *k* columns of the weight matrix (so that *W_j_*(*u*) = *Z_uj_*), set the (*k* + 1)^st^ column to the trait (so that *w*_*k*+1_(*u*) = *w_u_*), and the final column to all 1s (so *w*_*k*+2_(*u*) = 1). Also let *Z^T^h* = *v* be precomputed. Then the sum of the traits of samples with the focal genotype is *h_g_* = *x*_*k*+1_, and the allele count is *m* = *x*_*k*+2_, so that

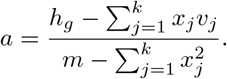

In practice we square this and divide by two, so that for biallelic loci the two alleles contribute an equal amount. For loci with more than two alleles, it would be more satisfying to return the proportion of variance in the trait that is explained by *all* of the alleles; however, this would be more involved (it would entail inversion of a 3 × 3 matrix for each locus).

## Supplementary figures

**Figure S1:**
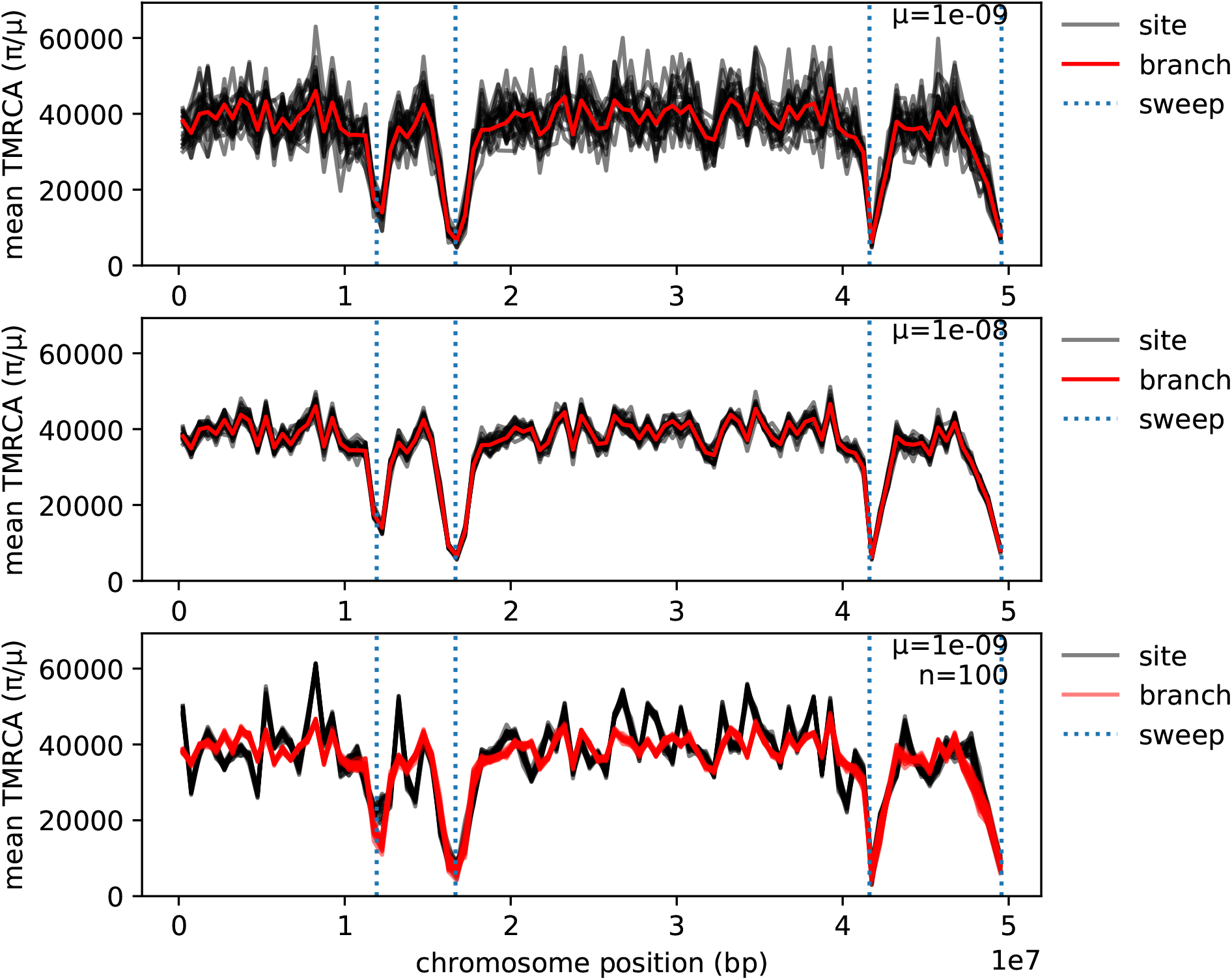
As in Figure 5, but in a population of size 10,000.

**Figure S2:**
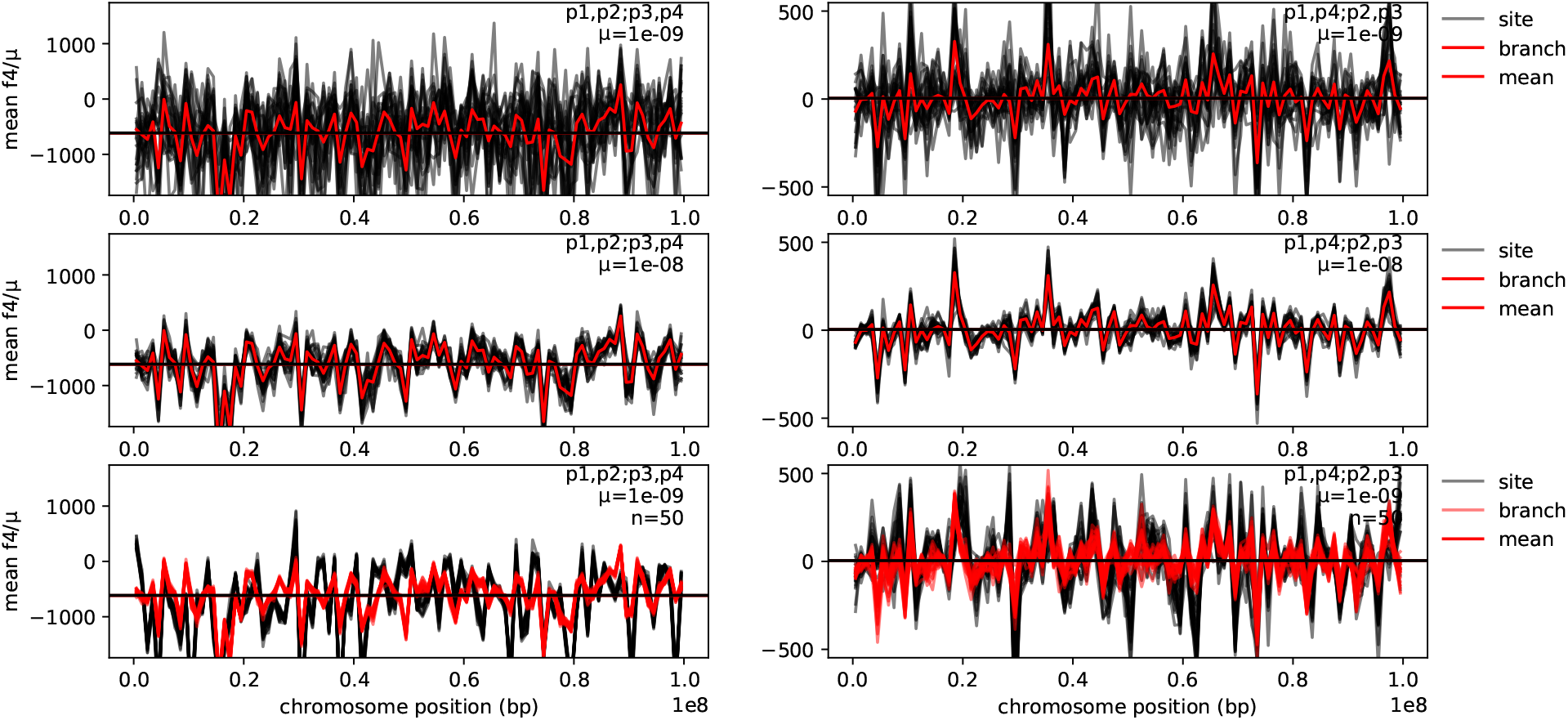
Values of the *f*_4_ statistic in megabase windows along the genome from the simulations of introgression described in the text. Statistics are shown in units of generations: Branch statistics are computed in units of time, which is in generations for these simulations; and Site statistics are reported in units of mutations per unit of sequence length, which we divide by mutation rate per unit of sequence length and per generation to obtain units of generations.

**Figure S3:**
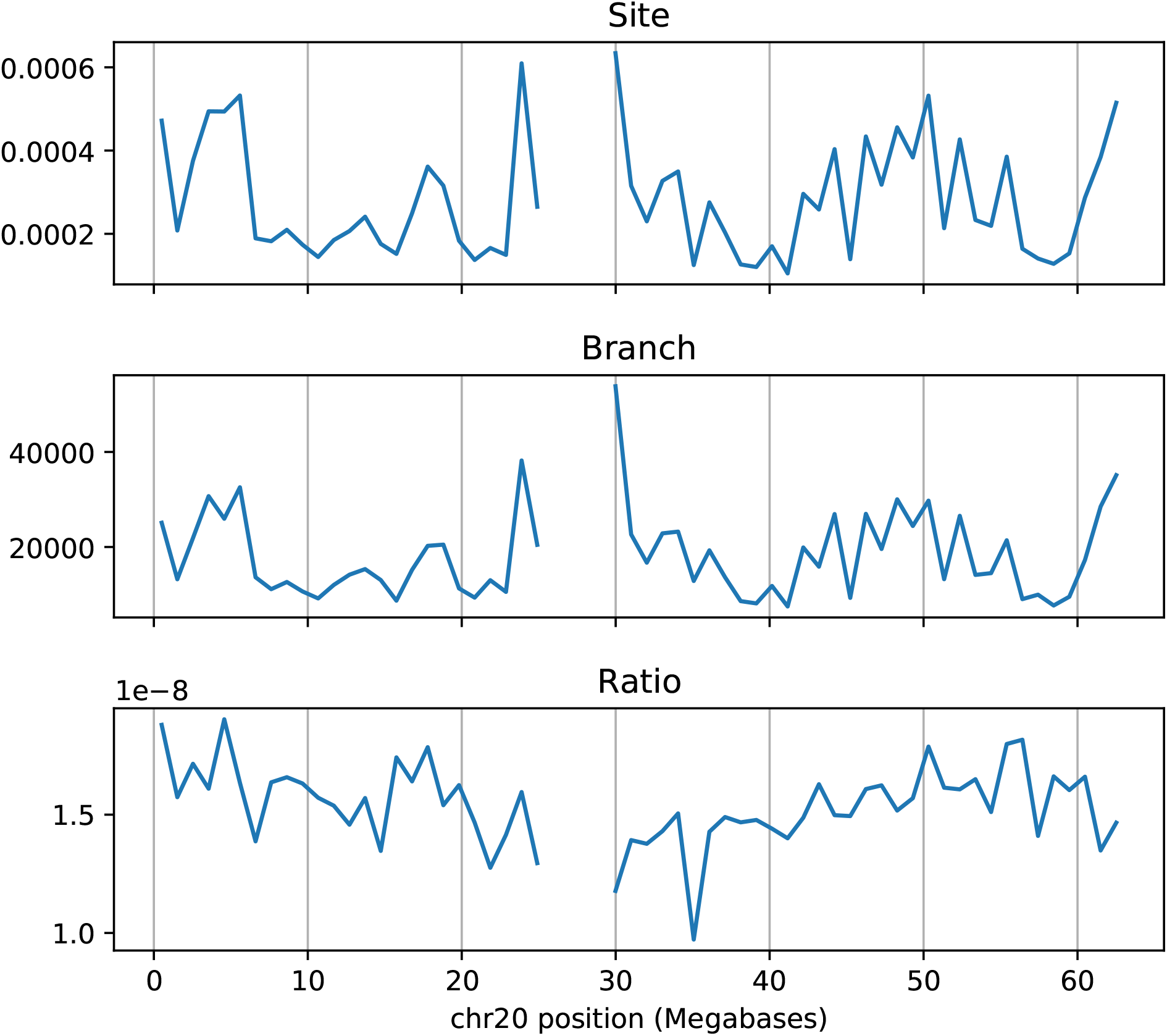
**(top)** Mean sequence diversity in 1Mb windows along human chromosome 20 of the GBR (Great Britain) sample of the 1000 Genomes Project, in units of mean number of pairwise differences per nucleotide. **(middle)** The dual Branch statistic as calculated from trees inferred by Relate [Speidel et al., 2019], in units of generations. **(bottom)** The ratio of the Site statistic to the Branch statistic. This figure differs from Figure 7 in that the trees were inferred using only GBR samples, and Relate was allowed to co-estimate population size and branch lengths, with mutation rate fixed at 1.45 × 10^-8^ mutations per generation; the trees shown in Figure 7 were inferred with a fixed effective population size of *N_e_* = 15, 000.

## Notes

https://github.com/petrelharp/treestats_ms

### Competing Interest Statement

The authors have declared no competing interest.

### Summary of Updates

We have added a few more explanatory figures and some additional discussion, mostly to the introductory section. We have also added more context for the Relate trees obtained from 1000 Genomes data.

## References

1000 Genomes Project Consortium. A global reference for human genetic variation. Nature, 526(7571):68–74, 2015.

Guillaume Achaz. Frequency spectrum neutrality tests: One for all and all for one. Genetics, 183(1):249–258, June 2009. doi: 10.1534/genetics.109.104042. URL https://doi.org/10.1534%2Fgenetics.109.104042.

Stepfanie M. Aguillon, John W. Fitzpatrick, Reed Bowman, Stephan J. Schoech, Andrew G. Clark, Graham Coop, and Nancy Chen. Deconstructing isolation-by-distance: The genomic consequences of limited dispersal. PLOS Genetics, 13(8):1–27, 08 2017. doi: 10.1371/journal.pgen.1006911. URL https://doi.org/10.1371/journal.pgen.1006911.

Cecile Ane and Michael J Sanderson. Missing the forest for the trees: Phylogenetic compression and its implications for inferring complex evolutionary histories. Systematic Biology, 54(1):146–157, 2005.

Mark A. Beaumont, W. Zhang, and D. J. Balding. Approximate Bayesian computation in population genetics. Genetics, 162:2025–2026, 2002.

Celine Becquet and Molly Przeworski. A new approach to estimate parameters of speciation models with application to apes. Genome research, 17(10):1505–1519, 2007.

Champak R Beeravolu, Michael J Hickerson, Laurent AF Frantz, and Konrad Lohse. ABLE: blockwise site frequency spectra for inferring complex population histories and recombination. Genome biology, 19(1): 145, 2018.

Tom R Booker and Peter D Keightley. Understanding the factors that shape patterns of nucleotide diversity in the house mouse genome. Molecular Biology and Evolution, page msy188, 2018. doi: 10.1093/molbev/msy188. URL http://dx.doi.org/10.1093/molbev/msy188.

Brian L Browning, Ying Zhou, and Sharon R Browning. A one-penny imputed genome from next-generation reference panels. The American Journal of Human Genetics, 2018.

S. R. Browning and B. L. Browning. High-resolution detection of identity by descent in unrelated individuals. Am. J. Hum. Genet., 86:526–539, April 2010.

Clare Bycroft, Colin Freeman, Desislava Petkova, Gavin Band, Lloyd T Elliott, Kevin Sharp, Allan Motyer, Damjan Vukcevic, Olivier Delaneau, Jared O’Connell, et al. The UK Biobank resource with deep phenotyping and genomic data. Nature, 562:203–209, 2018.

B Charlesworth, M Nordborg, and D Charlesworth. The effects of local selection, balanced polymorphism and background selection on equilibrium patterns of genetic diversity in subdivided populations. Genet Res, 70(2):155–174, October 1997. URL https://www.ncbi.nlm.nih.gov/pubmed/9449192.

Scott Christley, Yiming Lu, Chen Li, and Xiaohui Xie. Human genomes as email attachments. Bioinformatics, 25(2):274–275, 2008.

Katalin Csilléry, Michael GB Blum, Oscar E Gaggiotti, and Olivier François. Approximate Bayesian computation (ABC) in practice. Trends in Ecology & Evolution, 25(7):410–418, 2010.

Agnieszka Danek and Sebastian Deorowicz. GTC: how to maintain huge genotype collections in a compressed form. Bioinformatics, 34(11):1834–1840, 2018.

Richard Durbin. Efficient haplotype matching and storage using the positional Burrows-Wheeler transform (PBWT). Bioinformatics, 30(9):1266–1272, 2014.

David Eppstein. Offline algorithms for dynamic minimum spanning tree problems. Journal of Algorithms, 17(2):237 – 250, 1994. ISSN 0196-6774. doi: https://doi.org/10.1006/jagm.1994.1033. URL http://www.sciencedirect.com/science/article/pii/S0196677484710339.

David Eppstein, Zvi Galil, Giuseppe F. Italiano, and Amnon Nissenzweig. Sparsification - a technique for speeding up dynamic graph algorithms. J. ACM, 44(5):669–696, September 1997. ISSN 0004-5411. doi: 10.1145/265910.265914. URL http://doi.acm.org/10.1145/265910.265914.

Joseph Felsenstein. Inferring phylogenies. Sinauer associates Sunderland, MA, 2004.

Luca Ferretti, Alice Ledda, Thomas Wiehe, Guillaume Achaz, and Sebastian E. Ramos-Onsins. Decomposing the site frequency spectrum: The impact of tree topology on neutrality tests. Genetics, 207(1):229–240, 2017. ISSN 0016-6731. doi: 10.1534/genetics.116.188763. URL https://www.genetics.org/content/207/1/229.

Yun Xin Fu. Statistical properties of segregating sites. Theor Popul Biol, 48(2):172–197, October 1995. doi: 10.1006/tpbi.1995.1025. URL http://www.ncbi.nlm.nih.gov/pubmed/7482370.

Jared Galloway, William A Cresko, and Peter Ralph. A few stickleback suffice for the transport of alleles to new lakes. G3: Genes, Genomes, Genetics, 10(2):505–514, 2020.

John H. Gillespie and Charles H. Langley. Are evolutionary rates really variable? Journal of Molecular Evolution, 13(1):27–34, 1979. ISSN 0022-2844. doi: 10.1007/BF01732751. URL http://dx.doi.org/10.1007/BF01732751.

Robert C Griffiths. The two-locus ancestral graph. Lecture Notes-Monograph Series, pages 100–117, 1991.

Robert C Griffiths and Paul Marjoram. Ancestral inference from samples of DNA sequences with recombination. Journal of Computational Biology, 3(4):479–502, 1996.

Quiterie Haenel, Telma G. Laurentino, Marius Roesti, and Daniel Berner. Meta-analysis of chromosome-scale crossover rate variation in eukaryotes and its significance to evolutionary genomics. Molecular Ecology, 27(11):2477–2497, 2018. doi: 10.1111/mec.14699. URL https://onlinelibrary.wiley.com/doi/abs/10.1111/mec.14699.

Benjamin C Haller and Philipp W Messer. SLiM 3: forward genetic simulations beyond the Wright-Fisher model. Molecular biology and evolution, 36(3):632–637, 2019.

Benjamin C Haller, Jared Galloway, Jerome Kelleher, Philipp W Messer, and Peter L Ralph. Tree-sequence recording in SLiM opens new horizons for forward-time simulation of whole genomes. Molecular ecology resources, 2018.

Kelley Harris. From a database of genomes to a forest of evolutionary trees. Nature genetics, 51(9):1306–1307, 2019.

Jacob Holm, Kristian De Lichtenberg, Mikkel Thorup, and Mikkel Thorup. Poly-logarithmic deterministic fully-dynamic algorithms for connectivity, minimum spanning tree, 2-edge, and biconnectivity. Journal of the ACM (JACM), 48(4):723–760, 2001. URL http://doi.acm.org/10.1145/502090.502095.

Richard R Hudson. Properties of a neutral allele model with intragenic recombination. Theor Popul Biol, 23(2):183–201, April 1983. URL https://www.ncbi.nlm.nih.gov/pubmed/6612631.

Richard R Hudson. How can the low levels of DNA sequence variation in regions of the Drosophila genome with low recombination rates be explained? Proc Natl Acad Sci USA, 91(15):6815–6818, July 1994. URL https://www.ncbi.nlm.nih.gov/pubmed/8041702.

Richard R. Hudson. Two-locus sampling distributions and their application. Genetics, 159(4):1805–1817, 2001. ISSN 0016-6731. URL https://www.genetics.org/content/159/4/1805.

Konrad J Karczewski, Laurent C Francioli, Grace Tiao, Beryl B Cummings, Jessica Alföldi, Qingbo Wang, Ryan L Collins, Kristen M Laricchia, Andrea Ganna, Daniel P Birnbaum, et al. Variation across 141,456 human exomes and genomes reveals the spectrum of loss-of-function intolerance across human proteincoding genes. BioRxiv, page 531210, 2019.

Jerome Kelleher, Alison M Etheridge, and Gilean McVean. Efficient coalescent simulation and genealogical analysis for large sample sizes. PLoS computational biology, 12(5):e1004842, 2016.

Jerome Kelleher, Kevin R. Thornton, Jaime Ashander, and Peter L. Ralph. Efficient pedigree recording for fast population genetics simulation. PLoS Computational Biology, 14(11):1–21, 11 2018. doi: 10.1371/journal.pcbi.1006581. URL https://doi.org/10.1371/journal.pcbi.1006581.

Jerome Kelleher, Yan Wong, Anthony W. Wohns, Chaimaa Fadil, Patrick K. Albers, and Gil McVean. Inferring whole-genome histories in large population datasets. Nature Genetics, 51(9):1330–1338, 2019. ISSN 15461718. doi: 10.1038/s41588-019-0483-y. URL https://doi.org/10.1038/s41588-019-0483-y.

M Kreitman. Nucleotide polymorphism at the alcohol dehydrogenase locus of *Drosophila melanogaster*. Nature, 304(5925):412–417, 1983. ISSN 0028-0836. doi: 10.1038/304412a0. URL http://dx.doi.org/10.1038/304412a0.

Jakub Lacki and Piotr Sankowski. Reachability in graph timelines. In Proceedings of the 4th Conference on Innovations in Theoretical Computer Science, ITCS ‘13, pages 257–268, New York, NY, USA, 2013. ACM. ISBN 978-1-4503-1859-4. doi: 10.1145/2422436.2422468. URL http://doi.acm.org/10.1145/2422436.2422468.

Ryan M Layer, Neil Kindlon, Konrad J Karczewski, Aaron R Quinlan, Exome Aggregation Consortium, et al. Efficient genotype compression and analysis of large genetic-variation data sets. Nature methods, 13 (1):63, 2016.

Ellen M. Leffler, Kevin Bullaughey, Daniel R. Matute, Wynn K. Meyer, Laure Ségurel, Aarti Venkat, Peter Andolfatto, and Molly Przeworski. Revisiting an old riddle: What determines genetic diversity levels within species? PLoS Biol, 10(9):e1001388, 09 2012. doi: 10.1371/journal.pbio.1001388. URL http://dx.doi.org/10.1371%2Fjournal.pbio.1001388.

Haipeng Li and Thomas Wiehe. Coalescent tree imbalance and a simple test for selective sweeps based on microsatellite variation. PLoS Comput Biol, 9(5):e1003060, 2013.

Michael F Lin, Xiaodong Bai, William J Salerno, and Jeffrey G Reid. Sparse Project VCF: efficient encoding of population genotype matrices. BioRxiv, page 611954, 2019.

Konrad Lohse. Come on feel the noise-from metaphors to null models. J. Evol. Biol, 30:1506–1508, 2017.

Konrad Lohse, Martin Chmelik, Simon H Martin, and Nicholas H Barton. Efficient strategies for calculating blockwise likelihoods under the coalescent. Genetics, 202(2):775–786, 2016.

Alicia R Martin, Christopher R Gignoux, Raymond K Walters, Genevieve L Wojcik, Benjamin M Neale, Simon Gravel, Mark J Daly, Carlos D Bustamante, and Eimear E Kenny. Human demographic history impacts genetic risk prediction across diverse populations. The American Journal of Human Genetics, 100(4):635–649, 2017.

GAT McVean. A genealogical interpretation of linkage disequilibrium. Genetics, 162(2):987–991, October 2002. URL http://www.ncbi.nlm.nih.gov/pmc/articles/PMC1462283/.

Alistair Miles and Nick Harding. cggh/scikit-allel: v1.1.8, July 2017. URL https://doi.org/10.5281/zenodo.822784.

Mark J. Minichiello and Richard Durbin. Mapping trait loci by use of inferred ancestral recombination graphs. The American Journal of Human Genetics, 79(5):910 – 922, 2006. ISSN 0002-9297. doi: https://doi.org/10.1086/508901. URL http://www.sciencedirect.com/science/article/pii/S0002929707608349.

Vagheesh M Narasimhan, Raheleh Rahbari, Aylwyn Scally, Arthur Wuster, Dan Mason, Yali Xue, John Wright, Richard C Trembath, Eamonn R Maher, David A van Heel, et al. Estimating the human mutation rate from autozygous segments reveals population differences in human mutational processes. Nature communications, 8(1):303, 2017.

Nick Patterson, Priya Moorjani, Yontao Luo, Swapan Mallick, Nadin Rohland, Yiping Zhan, Teri Genschoreck, Teresa Webster, and David Reich. Ancient admixture in human history. Genetics, 192(3): 1065–1093, 2012.

Shaun Purcell, Benjamin Neale, Kathe Todd-Brown, Lori Thomas, Manuel AR Ferreira, David Bender, Julian Maller, Pamela Sklar, Paul IW De Bakker, Mark J Daly, et al. Plink: a tool set for whole-genome association and population-based linkage analyses. The American journal of human genetics, 81(3):559–575, 2007.

Dandi Qiao, Wai-Ki Yip, and Christoph Lange. Handling the data management needs of high-throughput sequencing data: Speedgene, a compression algorithm for the efficient storage of genetic data. BMC bioinformatics, 13(1):100, 2012.

Peter L. Ralph. An empirical approach to demographic inference with genomic data. Theoretical Population Biology, 127:91 – 101, 2019. ISSN 0040-5809. doi: https://doi.org/10.1016/j.tpb.2019.03.005. URL http://www.sciencedirect.com/science/article/pii/S0040580918301667.

Matthew D Rasmussen, Melissa J Hubisz, Ilan Gronau, and Adam Siepel. Genome-wide inference of ancestral recombination graphs. PLoS genetics, 10(5):e1004342, 2014.

David Reich, Kumarasamy Thangaraj, Nick Patterson, Alkes L Price, and Lalji Singh. Reconstructing indian population history. Nature, 461(7263):489, 2009.

Francesco Sambo, Barbara Di Camillo, Gianna Toffolo, and Claudio Cobelli. Compression and fast retrieval of SNP data. Bioinformatics, 30(21):3078–3085, 2014.

Christiana L. Scheib, Ruoyun Hui, Eugenia D’Atanasio, Anthony Wilder Wohns, Sarah A. Inskip, Alice Rose, Craig Cessford, Tamsin C. O’Connell, John E. Robb, Christopher Evans, Ricky Patten, and Toomas Kivisild. East Anglian early Neolithic monument burial linked to contemporary Megaliths. Annals of Human Biology, 46(2):145–149, 2019. doi: 10.1080/03014460.2019.1623912. URL https://doi.org/10.1080/03014460.2019.1623912.

Daniel R Schrider and Andrew D Kern. Supervised machine learning for population genetics: a new paradigm. Trends in Genetics, 34(4):301–312, 2018.

Charles Semple and Mike A Steel. Phylogenetics. Oxford University Press, 2003.

M Slatkin. Inbreeding coefficients and coalescence times. Genet Res, 58(2):167–175, October 1991. URL http://www.ncbi.nlm.nih.gov/pubmed/1765264.

Leo Speidel, Marie Forest, Sinan Shi, and Simon R. Myers. A method for genome-wide genealogy estimation for thousands of samples. Nature Genetics, 51(9):1321–1329, 2019. ISSN 15461718. doi: 10.1038/s41588-019-0484-x. URL https://doi.org/10.1038/s41588-019-0484-x.

S. Srinivasan, S. Pollard, S. K. Das, B. Norris, and S. Bhowmick. A shared-memory algorithm for updating tree-based properties of large dynamic networks. IEEE Transactions on Big Data, pages 1–1, 2018. doi: 10.1109/TBDATA.2018.2870136. URL https://ieeexplore.ieee.org/document/8464249.

Sean Stankowski, Madeline A. Chase, Allison M. Fuiten, Murillo F. Rodrigues, Peter L. Ralph, and Matthew A. Streisfeld. Widespread selection and gene flow shape the genomic landscape during a radiation of monkeyflowers. PLOS Biology, 17(7):1–31, 07 2019. doi: 10.1371/journal.pbio.3000391. URL https://doi.org/10.1371/journal.pbio.3000391.

Fran Supek and Ben Lehner. Differential DNA mismatch repair underlies mutation rate variation across the human genome. Nature, 521:81–, February 2015. URL https://doi.org/10.1038/nature14173.

F Tajima. Statistical method for testing the neutral mutation hypothesis by DNA polymorphism. Genetics, 123(3):585–595, November 1989. ISSN 0016-6731. URL https://www.ncbi.nlm.nih.gov/pubmed/2513255.

Fumio Tajima. Evolutionary relationship of DNA sequences in finite populations. Genetics, 105(2):437–460, 1983. URL https://www.genetics.org/content/105/2/437.long.

Simon Tavaré. Line-of-descent and genealogical processes, and their applications in population genetics models. Theoretical Population Biology, 26(2):119 – 164, 1984. ISSN 0040-5809. doi: 10.1016/0040-5809(84)90027-3. URL http://www.sciencedirect.com/science/article/pii/0040580984900273.

Kevin R Thornton. Libsequence: a C++ class library for evolutionary genetic analysis. Bioinformatics, 19(17):2325–2327, November 2003. ISSN 1367-4803. doi: 10.1093/bioinformatics/btg316. URL https://www.ncbi.nlm.nih.gov/pubmed/14630667.

Kevin R Thornton. A C++ template library for efficient forward-time population genetic simulation of large populations. Genetics, 198(1):157–166, 2014.

G A Watterson. On the number of segregating sites in genetical models without recombination. Theor. Popul. Biol., 7(2):256–276, April 1975. ISSN 0040-5809. doi: 10.1016/0040-5809(75)90020-9. URL https://www.ncbi.nlm.nih.gov/pubmed/1145509.

S Zöllner and J K Pritchard. Coalescent-based association mapping and fine mapping of complex trait loci. Genetics, 169(2):1071–1092, February 2005. doi: 10.1534/genetics.104.031799. URL https://www.ncbi.nlm.nih.gov/pmc/articles/PMC1449137/.

